# Conformations of a Low-Complexity Protein in Homogeneous and Phase-Separated Frozen Solutions

**DOI:** 10.1101/2024.07.25.605144

**Authors:** C. Blake Wilson, Myungwoon Lee, Wai-Ming Yau, Robert Tycko

**Author notes:** corresponding author: Dr. Robert Tycko, National Institutes of Health, Building 5, Room 409, Bethesda, MD 20892-0520.

## Abstract

Solutions of the intrinsically disordered, low-complexity domain of the FUS protein (FUS-LC) undergo liquid-liquid phase separation (LLPS) below temperatures T_LLPS_ in the 20-40° C range. To investigate whether local conformational distributions are detectably different in the homogeneous and phase-separated states of FUS-LC, we performed solid state nuclear magnetic resonance (ssNMR) measurements on solutions that were frozen on sub-millisecond time scales after equilibration at temperatures well above (50° C) or well below (4° C) T_LLPS_. Measurements were performed at 25 K with signal enhancements from dynamic nuclear polarization. Crosspeak patterns in two-dimensional (2D) ssNMR spectra of rapidly frozen solutions in which FUS-LC was uniformly ^15^N,^13^C-labeled were found to be nearly identical for the two states. Similar results were obtained for solutions in which FUS-LC was labeled only at Thr, Tyr, and Gly residues, as well as solutions of a FUS construct in which five specific residues were labeled by ligation of synthetic and recombinant fragments. These experiments show that local conformational distributions are nearly the same in the homogeneous and phase-separated solutions, despite the much greater protein concentrations and more abundant intermolecular interactions within phase-separated, protein-rich “droplets”. Comparison of the experimental results with simulations of the sensitivity of 2D crosspeak patterns to an enhanced population of β-strand-like conformations suggests that changes in conformational distributions are no larger than 5-10%.

**Statement of Significance:** Liquid-liquid phase separation (LLPS) in solutions of proteins with intrinsically disordered domains has attracted recent attention because of its relevance to multiple biological processes and its inherent interest from the standpoint of protein biophysics. The high protein concentrations and abundant intermolecular interactions within protein-rich, phase-separated “droplets” suggests that conformational distributions of intrinsically disordered proteins may differ in homogeneous and phase-separated solutions. To investigate whether detectable differences exist, we performed experiments on the low-complexity domain of the FUS protein (FUS-LC) in which FUS-LC solutions were first equilibrated at temperatures well above or well below their LLPS transition temperatures, then rapidly frozen and examined at very low temperatures by solid state nuclear magnetic resonance (ssNMR) spectroscopy. The ssNMR data for homogeneous and phase-separated frozen solutions of FUS-LC were found to be nearly identical, showing that LLPS is not accompanied by substantial changes in the local conformational distributions of this intrinsically disordered protein.

## Introduction

Liquid-liquid phase separation (LLPS) in biopolymer solutions is a topic of active research due to its biological relevance and inherent biophysical interest (1–5). While LLPS in protein solutions has been studied previously, for example in the context of crystallization mechanisms (6), recent studies have focus on LLPS in solutions of proteins that are intrinsically disordered or have intrinsically disordered domains (7–25). By definition, intrinsically disordered proteins (IDPs) lack persistent secondary and tertiary structure and have broad conformational distributions in their monomeric states, approximating the conformational distribution of a random coil. However, IDPs may become conformationally ordered upon binding to protein partners in a complex (26). IDPs also adopt well-defined structures when they self-assemble to form amyloid fibrils (27–34). Development of structural order in a complex or fibril depends on intermolecular interactions. The possibility therefore exists that the level of conformational order and the conformational preferences within an IDP may change when phase separation occurs, since both the local protein concentration and the fraction of time during which each protein chain engages in intermolecular interactions increase substantially in the protein-rich “droplet” regions of the phase-separated state.

It has been argued that intermolecular interactions between IDPs are similar to intramolecular interactions within an IDP, so that changes in conformational distributions upon phase separation should not be expected (14,22). As a possible counter-argument, it is worth noting that amyloid fibril formation by IDPs typically results in cross-β structures that contain in-register parallel β-sheets, i.e., β-sheets in which identical amino acid sequences from neighboring molecules align with one another (27–34). Such structures are adopted generically in amyloid fibrils because they permit the intermolecular backbone hydrogen bonds of a β-sheet to be combined with favorable intermolecular interactions of like sidechains (*e.g.*, contacts between hydrophobic sidechains, π-stacking of aromatic sidechains, hydrogen bonds among sidechains in rows of Gln or Asn residues). In addition, certain IDPs that exhibit LLPS also form amyloid fibrils at longer times (5,7,10,16–18,35), implying that the liquid-liquid phase-separated state is a metastable condition, with the fibrillar state being the true thermodynamic equilibrium state. For example, the intrinsically disordered, low-complexity (LC) domains of RNA-binding proteins such as FUS (FUsed in Sarcoma), heterologous nuclear ribonucleoproteins (hnRNPs), TDP-43, and TIA1 exhibit this behavior (7,10,15,17,18,28,35,36). If intermolecular interactions in the phase-separated state are related to intermolecular interactions in the fibrillar state, then one might expect conformational distributions of IDPs to develop greater populations of β-strand-like conformations when LLPS occurs, at least for certain sites or segments within the IDP sequence (37,38).

On the experimental side, information about the effect of LLPS on the conformational distributions of IDPs comes primarily from magnetic resonance spectroscopy. Nuclear magnetic resonance (NMR) studies of phase-separated protein solutions have been reported by a number of groups (8,9,11,19–21,39–49). These studies are subject to limitations imposed by the high viscosity within phase-separated droplets, which impairs the resolution of the solution NMR spectra and the sensitivity of multidimensional measurements that depend on long time periods for nuclear spin polarization transfers. Nonetheless, NMR techniques and measurement conditions that lead to informative data have been developed (20,40,42,47). Solution NMR measurements on IDPs in the phase-separated state show that, despite the greatly reduced translational diffusion rates within protein-rich droplets, large-amplitude local motions can remain sufficiently rapid to enable multidimensional spectroscopy. NMR chemical shifts generally remain similar to random-coil values, implying an absence of persistent secondary structure. Measurements that test for exchange between unstructured and partially structured states on time scales greater than microseconds generally fail to find evidence for such conformational exchange processes. On the other hand, chemical shifts in the homogeneous state are not identical to chemical shifts in the phase-separated state (9,21,43,50). In the case of LLPS by the tau protein, chemical shift changes do indicate an increase in β-strand populations in the phase-separated state (47). Solution NMR measurements on the measles virus N_TAIL_ protein (21) and the Ddx4 protein (51) do indicate exchange between populations with distinct structural or dynamical properties within protein-rich regions of the phase-separated state.

Electron paramagnetic resonance (EPR) methods have also been used to study effects of LLPS on the structural and dynamical properties of IDPs (11,52–54). For the LC domain of FUS, measurements of distance distributions between electron spin labels indicate an overall compaction of protein monomers in the phase-separated state relative to monomers in a homogeneous solution (11,54). An EPR study of LLPS by the intrinsically disordered N-terminal domain of CPEB4 found evidence for partitioning of protein states with distinct dynamical properties between protein-rich droplets and the surrounding dilute solution (53). In addition, solid state NMR (ssNMR) techniques have been used by several groups to characterize molecular structures of protein fibrils that develop from phase-separated solutions of IDPs and to follow the time-dependent development of solid-like or gel-like states after LLPS (7,10,15–18,28,55).

Conformational properties of IDPs in phase-separated states have also been examined by computational methods, using either coarse-grained (14,22) or all-atom (12,23,56) representations of IDPs. For the LC domain of FUS, all-atom simulations do not show enhanced population of β-strand conformations in the phase-separated state (56).

In contrast, biochemical studies provide evidence for a correspondence between structural properties of IDPs in phase-separated droplets and in cross-β fibrils (28,57–59). Specifically, patterns of protection from sidechain acetylation by N-acetylimidazole were found to be similar in droplets and fibrils formed by the LC domain of hnRNPA2 (57), and greatest protection occurred in the C-terminal segment identified as the fibril core in subsequent structural studies by ssNMR (60) and cryo-electron microscopy (61). Site-directed mutagenesis experiments suggested that droplets formed by the LC domain of FUS are most readily susceptible to dissolution by enzymatic phosphorylation at Ser and Thr sites that lie within the fibril core-forming segment (28). For the LC domain of TDP-43, patterns of protection from Met sidechain oxidation by H_2_O_2_ were found to be similar in droplets and fibrils (58). Site-specific introduction of N-methyl amino acids, to eliminate the possibility of intermolecular backbone hydrogen bonding, was found to increase the susceptibility of droplets to dissolution by urea within the same segment of the TDP-43 LC domain that was most strongly protected from Met sidechain oxidation (59).

Thus, although the conformational state of IDPs in phase-separated droplets has been investigated previously, information from new experimental approaches may further refine our understanding of LLPS at the molecular structural level. This paper reports the results of experiments in which ssNMR measurements on rapidly frozen solutions were used to compare the conformational properties of an IDP in homogenous and phase-separated states. Experiments were performed on the LC domain of FUS, using the constructs shown in Figs. 1a and 1b (heretofore called FUS-LC and FUS_61-214_, respectively) with various isotopic labeling patterns. Samples for ssNMR were prepared by rapid freezing from temperatures well above or well below the phase transition temperature T_LLPS_. Two-dimensional (2D) ssNMR spectra of FUS-LC that was uniformly ^15^N,^13^C-labeled (U-^15^N,^13^C-FUS-LC) show only minor differences in ^13^C-^13^C crosspeak patterns. Similar results were obtained for frozen solutions of FUS-LC that was ^15^N,^13^C-labeled only at Thr, Tyr, and Gly residues (TYG-^15^N,^13^C-FUS-LC) and for frozen solutions of FUS_61-214_ that was selectively labeled only at five specific residues (sel-^15^N,^13^C-FUS_61-214_). Since NMR chemical shifts are sensitive to local conformations, our data provide evidence that major changes in conformational distributions do not occur when FUS-LC solutions undergo LLPS. From simulations of the effects of changes in conformational distributions on 2D ssNMR lineshapes, we estimate that populations of β-strand-like conformations in the phase-separated state may differ from those in the homogeneous state by no more than 5-10%.

**Figure 1:**
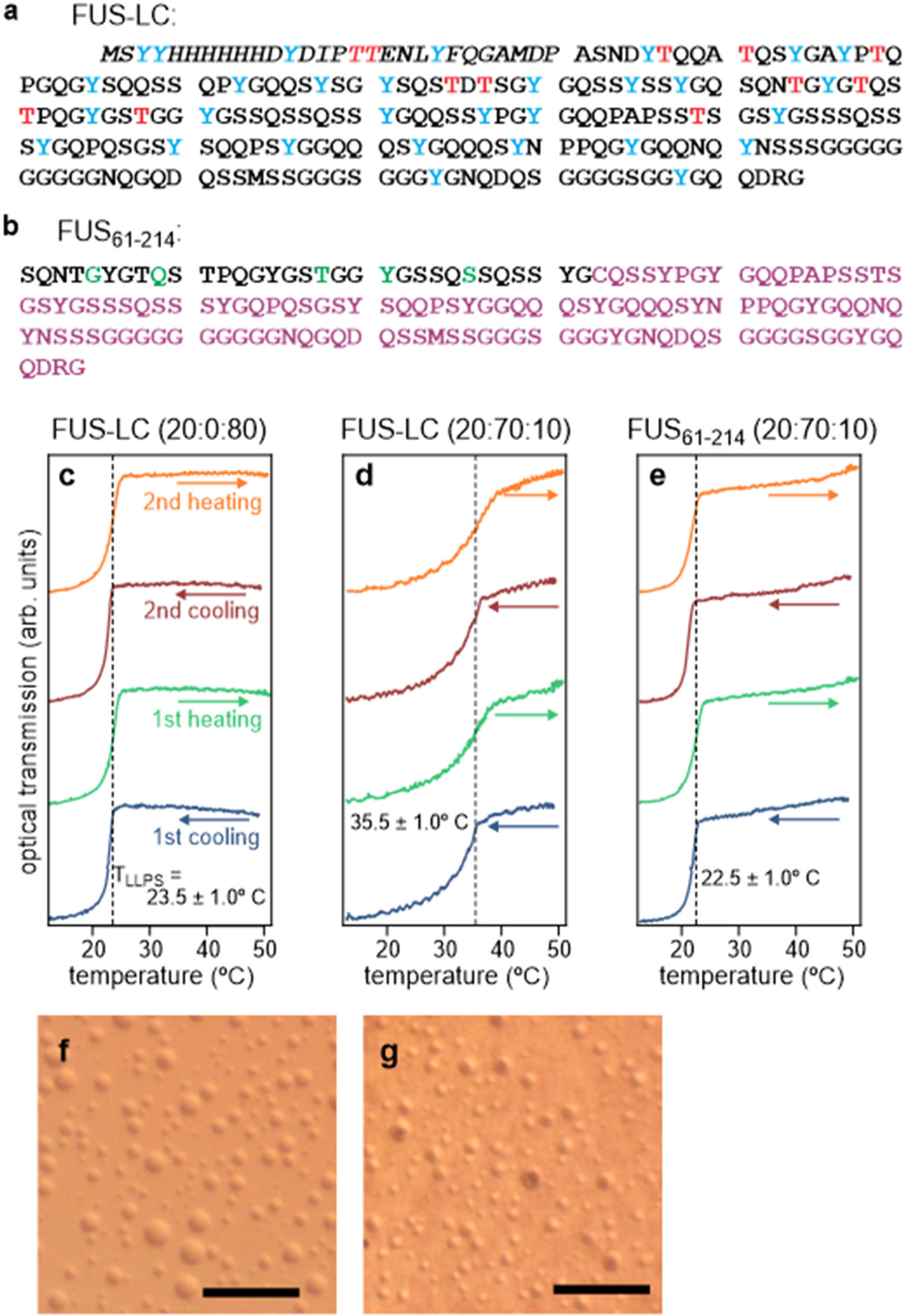
Characterization of LLPS by the LC domain of FUS. (a) Amino acid sequence of the FUS-LC construct used in ssNMR measurements. Italicized segment is an N-terminal purification tag, which was not removed. Thr and Tyr residues, which were isotopically labeled in TYG-^15^N,^13^C-FUS-LC samples, are highlighted in red and blue, respectively. (b) Sequence of the FUS_61-214_ construct, comprising a synthetic segment for residues 61-92, with five ^15^N,^13^C-labeled residues shown in green, and a recombinant segment for residues 93-214. A Gln-to-Cys substitution at residue 93 enabled ligation of the two segments. (c) Temperature dependence of the turbidity of a 5.0 mg/mL solution of FUS-LC through two cycles of cooling and heating between 50° C and 10° C. The solvent was 20 mM sodium phosphate, 50 mM NaCl, and 180 mM GdmCl, pH 7.4, in a 20:0:80 volume ratio of perdeuterated glycerol, D_2_O, and H_2_O. These data indicate a phase transition temperature T_LLPS_ = 23.5 ± 1.0° C. (d) Same as panel c, but with a 20:70:10 volume ratio of perdeuterated glycerol, D_2_O, and H_2_O, indicating T_LLPS_ = 35.5 ± 1.0° C. (e) Same as panel d, but for a solution containing 14 mg/ml of FUS_61-214_, indicating T_LLPS_ = 22.5 ± 1.0° C. (f) Optical microscope image of phase-separated FUS-LC droplets at 24° C. The solution was prepared with 5.0 mg/mL FUS-LC, 20 mM sodium phosphate, 50 mM NaCl, and 0.6 M GdmCl, in a 20:70:10 volume ratio of perdeuterated glycerol, D_2_O, and H_2_O. (g) Optical microscope image of the same sample at approximately 100 K. The sample was contained between sapphire plates and was frozen by immersion in liquid nitrogen. Scale bars are 100 μm.

## Materials and Methods

### Expression and purification of full-length FUS-LC

Recombinant proteins were expressed in BL21 (DE3) *Escherichia coli* cells. As previously described (62), full-length FUS-LC (residues 2-214 of human FUS, UniProt entry P35637-1) was expressed with a N-terminal His_6_ tag that includes a flexible linker and a tobacco etch virus (TEV) cleavage site. The amino acid sequence is shown in Fig. 1a. The N-terminal tag was left in place for the experiments described here. For all expressions, ampicillin was added to 100 μg/mL. Unlabeled FUS-LC was expressed in cells grown in Luria-Bertani (LB) medium. The cells were incubated in 10 mL of LB medium, shaken at 37° C overnight, then added to 990 mL of LB medium and shaken at 37° C until the optical density at 600 nm (OD_600_) reached 0.8-1.0 (4-5 h). The 1.0 L culture was transferred to a 25° C shaking incubator for 30 min before adding isopropyl β-D-1-thiogalactopyranoside (IPTG) to a final concentration of 0.5 mM to induce FUS-LC overexpression. The 1.0 L culture was then shaken for 20 h at 25° C, whereupon the cells were harvested by centrifugation and the pellet was frozen and stored at -80° C.

Uniformly labeled FUS-LC (U-^15^N,^13^C-FUS-LC) was expressed in M9 medium using 2 g/L of ^13^C-labeled glucose and 1 g/L of ^15^NH_4_Cl. Cells were incubated in 10 mL of LB medium with shaking at 37° C for 4 h, then pelleted and resuspended in 10 mL of M9 medium and shaken at 37° C overnight. The 10 mL overnight culture was added to 990 mL of M9 medium and shaken at 37° C until OD_600_ reached 0.8. The 1.0 L culture was transferred to a 25° C shaking incubator for 30 min before adding IPTG to a final concentration of 0.5 mM and was then shaken at 25° C for 20 h. Finally, the cells were harvested by centrifugation and the pellet was frozen and stored at -80° C.

FUS-LC with ^15^N,^13^C-labeled Thr and Tyr was expressed in a defined amino acid medium (DAM) consisting of M9 medium with the addition of 100 mg/L of ^15^N,^13^C-labeled L-tyrosine, 90 mg/L of ^15^N,^13^C-labeled L-threonine, 200 mg/L of unlabeled L-histidine, L-phenylalanine, L-proline, and L-tryptophan, and 100 mg/L of all other amino acids (unlabeled). All amino acids were added to the medium prior to autoclaving, and 2.0 g/L of unlabeled glucose, 1.0 g/L ^14^NH_4_Cl, and 1.0 mL of 100 mg/ml ampicillin were added through a 0.2 μm syringe filter after autoclaving. Cells were incubated in 10 mL of LB medium and shaken at 37° C for 1 h, then pelleted and resuspended in 10 mL of DAM and shaken at 37° C overnight. The 10 mL culture was added to 990 mL of DAM and shaken at 37° C until OD_600_ reached 0.8, when it was transferred to a 25° C shaking incubator for 30 min before adding IPTG to a final concentration of 0.5 mM. The culture was shaken at 25° C for 18 h, the cells were harvested by centrifugation, and the pellet was frozen and stored at -80° C. As discussed below, ssNMR spectra indicated that Gly residues were partially ^15^N,^13^C-labeled in this sample, in addition to the deliberate labeling of Thr and Tyr residues. We therefore call this sample TYG-^15^N,^13^C-FUS-LC.

U-^15^N,^13^C-FUS-LC and TYG-^15^N,^13^C-FUS-LC were purified in denaturing conditions as follows. Cell pellets were resuspended in 50 mL lysis buffer containing 50 mM Tris-HCl at pH 7.5, 500 mM NaCl, 20 mM β-mercaptoethanol (BME), 1% v/v Triton X-100, 1 tablet of SIGMAFAST EDTA-free protease inhibitor cocktail (Sigma-Aldrich), 0.2 mg/mL lysozyme, and 2.0 M guanidinium chloride (GdmCl), incubated on ice for 30 min, then sonicated in an ice bath using a Branson Model 250 sonifier with a tapered 1/4” tip at 0.5 output, 50% duty cycle for 10 min. Cell lysates were centrifuged at 140000 × g for 60 min at 4° C. The supernatant was mixed with 25 mL of NiNTA resin and rotated at 4° C overnight. The NiNTA resin was then packed into a gravity flow column (Thermo Scientific disposable polypropylene) and washed with 120 mL of buffer containing 20 mM Tris-HCl, pH 7.5, 500 mM NaCl, 20 mM BME, 0.1 mM phenylmethylsulfonyl fluoride (PMSF), 2.0 M GdmCl, and 20 mM imidazole. All buffers used during protein purification were filtered with a 0.22 μm bottle-top vacuum filter (Corning) or with a 0.22 μm Millex syringe filter (Millipore). His-tagged proteins were eluted using the same buffer, but with 250 mM imidazole. After elution, ethylenediamine tetraacetic acid (EDTA) was added to 0.5 mM.

U-^15^N,^13^C-FUS-LC and TYG-^15^N,^13^C-FUS-LC were further purified and buffer-exchanged with fast protein liquid chromatography (FPLC), using a HiLoad Superdex 200 pg column with 20 mM sodium phosphate pH 7.4, 500 mM NaCl, 20 mM BME, 0.1 mM PMSF, and 2.0 M GdmCl. After collection, protein samples were concentrated to 10 mg/mL using Amicon Ultra centrifugal filters with a 10 kDa cutoff (Millipore), with 6.0 M GdmCl. Samples were aliquoted into 0.5 ml tubes, flash frozen in liquid nitrogen, and stored at -80° C.

### Production of sel-^15^N,^13^C-FUS_61-214_ by ligation of synthetic and recombinant fragments

FUS_61-214_ was prepared by ligation of a synthetic peptide, representing residues 61-92 of FUS-LC (FUS_61-92_), with a recombinant peptide, representing residues 93-214 with a Gln-to-Cys substitution at residue 93 (Q93C-FUS-LC_93-214_). This semisynthetic approach (63,64) allowed ^15^N,^13^C-labeled amino acids to be introduced selectively at five specific sites, namely G65, Q69, T78, Y81, and S86. We call the resulting sample sel-^15^N,^13^C-FUS_61-214_. The amino acid sequence is shown in Fig. 1b.

FUS-LC_61-92_-NHNH_2_ was synthesized with the five labeled residues on a nominal 0.1 mmol scale on a Biotage Initiator+ Alstra solid phase peptide synthesizer, using fluorenylmethyloxycarbonyl (Fmoc) chemistry and activation with N,N’-diisopropylcarbodiimide (DIC) and OxymaPure (EMD Millipore) in N-methyl-2-pyrrolidone. As described by Zheng *et al*. (65), 2-chlorotrityl chloride resin (1.18 meq/g AAAPTec) was activated by refluxing with thionyl chloride (1.2 eq, Sigma-Aldrich) and pyridine (2.5 eq, Sigma-Aldrich) in dichloromethane for 4 h before hydrazination with hydrazine hydrate (50-60%, Sigma-Aldrich). Unlabeled residues were double-coupled with a five-fold excess of Fmoc-amino for 10 min at 75° C. For isotopically labeled residues, a three-fold excess of the labeled Fmoc-amino acid was single-coupled for 20 min at 75° C, followed by single-coupling with a five-fold excess of unlabeled Fmoc-amino acid for 5 min at 75° C. Deprotection with 20% piperidine (Sigma-Aldrich) and 0.1 M 1-hydroxybenzotriazole hydrate (HOBt, Chem-Impex Int’l Inc) in dimethylformamide was performed at 70° C for 5 min. Cleavage of the synthesis resin was performed with a standard cocktail (trifluoroacetic acid/triisopropylsilane/water, 9.5/2.5/2.5 mL) for 2.0 h. After filtering and washing the resin with trifluoroacetic acid, the crude product was precipitated by cold methyl tert-butyl ether and the pellet was further washed with cold methyl tert-butyl ether twice. The final product was purified by preparative high-performance liquid chromatography (HPLC) on a Zorbax 300SBC3 reverse-phase column (Agilent Technologies).

Recombinant GFP-tagged Q93C-FUS-LC_93-214_ was expressed in BL21 (DE3) *Escherichia coli* cells. The cells were grown in 1.0 L of M9 medium containing 2.0 g of ^13^C_6_-glucose and 1.0 g of ^15^N-ammonium chloride at 37° C with shaking at 240 rpm until OD_600_ reached 0.8-1.0. Protein expression was induced by adding 0.5 mM IPTG and the cells were further cultured at 20° C overnight. After protein expression, the cells were centrifuged at 4,000 × g for 30 min at 4° C. The cell pellet was resuspended in 50 mL of lysis buffer (20 mM Tris-HCl, pH 7.4, 500 mM NaCl, 1.0 % v/v Triton X-100, 20 mM BME, one tablet of protease inhibitor cocktail, 2.0 M urea, and 0.2 mg/mL lysozyme) and sonicated in an ice bath for 10 min using a Branson Model 250 sonifier at output level 8 and 20% duty cycle, with a 0.5 inch horn tip. The lysed cells were centrifuged at 35,000 rpm for 1 h at 4° C. The supernatant was loaded onto a gravity flow column containing 15 mL of Ni-NTA agarose resin (Gold Bio) and the column was placed in a rotator for 30 min at 4° C to allow protein binding. The column was then washed with 250 mL of wash buffer (20 mM Tris-HCl, pH 7.4, 500 mM NaCl, 20 mM BME, 0.1 mM PMSF, 2.0 M urea, and 20 mM imidazole) and eluted with 50 mL of elution buffer (20 mM Tris-HCl, pH 7.4, 200 mM NaCl, 20 mM BME, 0.1 mM PMSF, 2.0 M urea, and 250 mM imidazole). 0.5 mM EDTA was added to the eluted protein solutions.

Purified GFP-tagged Q93C-FUS-LC_93-214_ was concentrated to approximately 70 mg/mL using an Amicon ultracentrifuge filter with a molecular weight cutoff of 10 kDa. The green fluorescent protein (GFP) tag was cleaved by diluting the protein to 2 mg/mL with a cleavage buffer (20 mM Tris-HCl, pH 7.4, 200 mM NaCl, 20 mM BME, 0.1 mM PMSF, and 5 mM DTT), followed by addition of caspase-3 at a 1:1000 mass ratio of caspase-3 to Q93C-FUS-LC_93-214_. The resulting solution was rotated at room temperature overnight. The solution was centrifuged to pellet the cleaved Q93C-FUS-LC_93-214_, and the supernatant was discarded. The pelleted material was dissolved in 1-2 ml of buffer containing 20 mM Tris-HCl, pH 7.4, 200 mM NaCl, and 5.0 M guanidine thiocyanate, and heated to 80° C for 10 min. Q93C-FUS-LC_93-214_ was purified by FPLC using a Superdex 200 pg column with 20 mM Tris-HCl, pH 7.4, 200 mM NaCl, 20 mM BME, 0.1 mM PMSF, and 2.0 M GdmCl for 150 min at a flow rate of 2.0 mL/min. Q93C-FUS-LC_93-214_ eluted between 110 and 130 min. Purified protein was concentrated to approximately 60 mg/mL in 6.0 M GdmCl using an Amicon ultracentrifuge filter with a molecular weight cutoff of 3 kDa and stored at -80° C.

For native chemical ligation, 10 mg of FUS_61-92_-NHNH_2_ was activated by 0.2 M KNO_2_ (Acros Organics) in 2.0 mL of 0.2 M phosphate solution containing 6.0 M GdmCl (pH 3.0-3.1) for 20 min at -15° C to form FUS_61-92_-N_3_, which was immediately mixed with 27 mg of recombinant Q93C-FUS-LC_93-214_ in 1.0 mL of 0.2 M phosphate solution containing 0.5 M 4-mercaptophenylacetic acid and 6.0 M GdmCl (pH 7.0-7.2). After adjusting to a final pH of 6.8-7.0, the ligation solution was stirred overnight at room temperature. Ligation was quenched by adding 0.4 mL of 0.1 M tris(2-carboxyethyl)phosphine in H_2_O. The resulting solution was stirred for additional 30 min before purification by HPLC. The final ligated product was obtained in 50-70% yield.

### Preparation of solutions for ssNMR

Full-length FUS-LC samples were further concentrated to 50 mg/mL in a buffer containing 20 mM sodium phosphate, 50 mM NaCl, 0.1 mM PMSF, and 6.0 M GdmCl, then diluted 10-fold to 20 mM sodium phosphate, 20 mM NaCl, 0.1 mM PMSF, 0.6 M GdmCl. Under these conditions, LLPS occurred near room temperature, as evidenced by increased sample turbidity (see below). Samples were incubated at 4° C for 2 min, after which the protein-rich droplets were spun down at 8000 rpm for 5 min. The supernatant was removed. The protein-rich phase, which formed a gel with 1-2% of the original volume of the solution, was resuspended in a buffer that was pre-heated to 50° C. The resulting solution contained 5.0 mg/mL FUS-LC, 20 mM sodium phosphate, 20 mM NaCl, 0.1 mM PMSF, 180 mM GdmCl, with 20% glycerol, 70% D_2_O, and 10 % H_2_O by volume. For NMR samples, ^12^C-glycerol-d_8_ (Cambridge Isotope Laboratories) was used, along with 10 mM sulfoacetyl-DOTOPA (66) for U-^15^N,^13^C-FUS-LC or 5.5 mM sulfoacetyl-DOTOPA for TYG-^15^N,^13^C-FUS-LC and sel-^15^N,^13^C-FUS_61-214_. Samples were heated to 50° C in a water bath, allowed to cool to room temperature, and re-heated to 50° C to ensure that droplets completely dissolved at elevated temperatures. Protein concentrations were measured using the optical absorbance at 280 nm as reported by a NanoDrop 2000c Spectrophotometer (Thermo Scientific), after 30-fold dilution in a buffer that contained 2.0 M GdmCl.

For experiments on sel-^15^N,^13^C-FUS_61-214_, lyophilized protein was dissolved directly in a buffer containing 20 mM sodium phosphate, 10 mM NaCl, 0.1 mM PMSF, and 2.0 mM tris(2-carboxyethyl)phosphine (TCEP), with 20% ^12^C-glycerol-d_8_, 70% D_2_O, and 10% H_2_O. The sel-^15^N,^13^C-FUS_61-214_ concentration was 14 mg/mL.

### Measurements of transition temperatures

LLPS transition temperatures T_LLPS_ were quantified with the home-built apparatus depicted in Fig. S1. This apparatus allows the temperature-dependent turbidities of multiple samples (20-30 μL each) to be tracked simultaneously while sample temperatures are ramped repeatedly between upper and lower values and at rates that can be controlled in software. Major components of the device include a CP-063HT thermoelectric cooler and TC-720 temperature controller (TE Technology, Inc.), an aluminum plate with sample wells that mounts on the thermoelectric cooler, circuitry to drive a 570 nm light-emitting diode (Rohm Semiconductor SLA-370MT3F) below and a phototransistor (Vishay Semiconductors TEPT4400) above each sample well, and a multi-channel analog-to-digital converter (DATAQ Instruments model DI-710-UH) to record the temperature of the aluminum plate and the light intensity that passes through each sample well as functions of time.

For measurements in Fig. 1, 25 μl volumes of FUS-LC or FUS_61-214_ were loaded into the sample wells, which were sealed on the bottom with transparent tape. A layer of mineral oil was placed on top of each protein solution to prevent evaporation. The aluminum plate temperature was ramped linearly from 50° C to 10° C and from 10° C to 50° C at 4.3 °/min. Between each cooling/heating cycle, the plate remained at 50° C for 6 min.

### Rapid freezing of FUS-LC solutions for ssNMR

Frozen FUS-LC solutions for ssNMR measurements were prepared using the rapid freezing apparatus depicted in Fig. S2, which is based on a previously-described design (67,68). The apparatus consists of a 26 cm long stainless steel tube with a 1 mm inner diameter soldered to a copper plate, to which a ceramic insulated strip heater (Omega Engineering, Inc. HSC-055-120V) is also attached. The temperature of the copper plate is monitored with a platinum resistor (Omega Engineering, Inc.). The stainless steel tube ends in a zero-dead-volume HPLC fitting (Valco Instruments) to which is attached a 1.0 cm length of 50 μm inner diameter PEEKSil HPLC tubing that acts as a nozzle. An HPLC pump (Teledyne ReaXus LS class) is connected to the upstream end of the stainless steel tubing. Pressure from the pump (900 psi at 4° C, 500 psi at 50° C) expels solutions from the stainless steel tube as high-speed jets.

The apparatus can hold up to 200 μL of liquid at a constant temperature. For each ssNMR sample, approximately 125 uL of FUS-LC solution was loaded into the apparatus with a syringe. The remaining volume was filled with hexane. The temperature of the apparatus was set either to 50° C (well above T_LLPS_) using the attached heater or to 4° C (well below T_LLPS_) by placing the entire apparatus in a cold room. Samples were incubated in the apparatus for 2.0 min to ensure equilibration, then expelled with a jet velocity of 8.5 m/s into a rapidly stirred isopentane bath, which was precooled to -145° C with liquid nitrogen. This protocol freezes samples in approximately 150 μs (68). Frozen sample particles were collected and packed into pre-cooled zirconia NMR rotors (46 mm length, 4 mm outer diameter, O’Keefe Ceramics) as previously described (67,68). Samples were stored in liquid nitrogen until ssNMR measurements were carried out.

Supernatant concentrations of FUS-LC or FUS_61-214_ solutions were less than 1 μM after incubation at 4° C and pelleting of phase-separated droplets. Therefore, less than 1% of the signal intensity in ssNMR spectra of samples that were rapidly frozen from the phase-separated state is attributable to protein molecules outside the droplets.

### ssNMR measurements

Measurements were carried out with a 9.4 T magnet (Oxford Instruments), a Bruker Avance III spectrometer operating at a ^1^H Larmor frequency of 400.8 MHz, and a home-built ssNMR probe with magic-angle spinning (MAS) and dynamic nuclear polarization (DNP) capabilities that has been described previously (69,70). The probe uses room-temperature nitrogen gas for magic-angle spinning (MAS) drive and air bearings and cryogenic helium (or nitrogen) gas to cool samples in the central region of the long NMR rotors. With helium cooling, sample temperatures were 25 K, as measured by ^79^Br spin-lattice relaxation times in a KBr powder (71). Liquid helium consumption was typically 1.8 L/h (70). The probe can be lowered beneath the NMR magnet while remaining cold and is designed to allow cold samples to be changed quickly (within 10 s). Sample temperatures remained below 100 K throughout the sample loading process.

Cross-effect DNP for ssNMR signal enhancements was driven by continuous-wave (CW) microwave irradiation at 263.9 GHz, near the low-frequency edge of absorption by unpaired electron spins of the sulfoacetyl-DOTOPA dopant (66). For measurements on U-^15^N,^13^C-FUS-LC and TYG-^15^N,^13^C-FUS-LC, microwaves were produced by a 1.5 W extended interaction oscillator (EIO, Communications & Power Industries). For measurements on sel-^15^N,^13^C-FUS_61-214_, the 1.5 W microwave source was unavailable and a 200 mW solid-state microwave source (Virginia Diodes, Inc) was used instead. A quasioptical interferometer (Thomas Keating, Ltd.) was used in all cases to produce circularly polarized microwaves, thereby maximizing the effective microwave power for DNP as previously described (70).

One-dimensional (1D) and 2D ssNMR spectra were acquired using pulse sequences shown in Fig. S3. At the start of each experiment, a train of π/2 pulses with separations τ_d_ was applied to all channels to destroy pre-existing nuclear spin polarizations. The ^1^H polarization was subsequently allowed to build up under DNP during a period τ_DNP_, with a time constant T_DNP_ that was measured for each sample. After the DNP build-up step, cross-polarization (CP) was used to transfer enhanced ^1^H polarization to ^13^C spins. DNP enhancement factors ε_DNP_, defined as the ratios of cross-polarized ^13^C ssNMR signal amplitudes with microwaves applied during τ_DNP_ to signal amplitudes without microwaves, were in the 45-55 range for rapidly frozen FUS-LC solutions at 25 K when the 1.5 W EIO source was used, and in the 20-30 range when the 200 mW solid-state microwave source was used. Values of ε_DNP_ and T_DNP_ are listed in Table S1. Fig. S4 shows examples of ssNMR data from which these values were determined.

Measurements on U-^15^N,^13^C-FUS-LC were carried out at a MAS frequency of 7.0 kHz, using radio-frequency (RF) amplitudes of 59 kHz on ^1^H and 52 kHz on ^13^C during CP. Lengths of π/2 pulses were 5.0 μs for ^1^H and ^13^C. Measurements on TYG-^15^N,^13^C-FUS-LC used a MAS frequency of 8.0 kHz, with RF amplitudes during CP of 60 kHz and 52 kHz on ^1^H and ^13^C, respectively. Measurements on sel-^15^N,^13^C-FUS_61-214_ used a MAS spinning speed of 8.0 kHz, with RF amplitudes during CP of 53 kHz and 37 kHz on ^1^H and ^13^C, respectively. Lengths of π/2 pulses were 5.0 μs for ^1^H and 7.0 μs for ^13^C. Two-pulse phase-modulated (TPPM) ^1^H decoupling (72) was applied with an 85 kHz RF amplitude during direct (t_2_) and indirect (t_1_) evolution periods. 2D ^13^C-^13^C measurements used maximum t_1_ values of 4 ms, spin diffusion mixing periods τ_SD_ equal to 20 ms, and total measurement times of 0.75-10.2 h. 2D ^15^N-^13^C measurements on TYG-^15^N,^13^C-FUS-LC use maximum t_1_ values of 3.5 ms, τ_SD_ = 25 ms, and total measurement times of 4.3 h. ^1^H-^15^N CP used RF amplitudes of 52 kHz on ^1^H and 36 kHz on ^15^N. ^15^N-^13^C CP used RF amplitudes of 11 kHz on ^15^N and 19 kHz on ^13^C, with the ^13^C RF carrier frequency set near the NMR frequencies of ^13^C_α_ sites for selectivity (73). ^15^N π/2 pulse lengths were 7 μs.

### ssNMR data analyses

Data analyses were carried out with Python scripts that made use of the NMRglue library (74). 1D slices through 2D spectra were extracted along the direct dimension and along the corresponding indirect dimension, where signals are expected to be symmetric about the diagonal in 2D ^13^C-^13^C experiments. 1D slices are plotted with the 50° C slices (*i.e.*, slices from spectra of samples that were rapidly frozen from 50° C) scaled to minimize the root-mean-squared (RMS) difference between the 50° C and 4° C slices. A 30 ppm region around the diagonal was excluded from RMS difference calculations. Difference spectra were calculated using these scaled slices. Center-of-mass calculations were carried out by taking the NMR signal *f* ( *x_i_* , *y _j_* ) in a rectangular *N* by *M* region, and calculating the center-of-mass coordinates (*x_c_* , *y_c_* ) according to 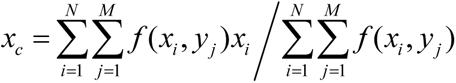 and 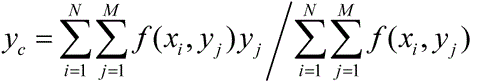. Uncertainties in the center-of-mass coordinates were estimated by repeating the calculation 128 times, each time adding noise to the region of interest with an RMS value equal to the RMS noise in the experimental 2D spectrum. Uncertainties are taken to be the standard deviations of the resulting distributions of center-of-mass coordinates.

### Optical microscopy at low temperatures

Experiments to visualize the effects of freezing on phase-separated solutions were carried out with the home-built cryogenic microscopy platform shown in Fig. S5. For these experiments, a solution of full-length FUS-LC (without isotopic labeling) was prepared as described above for ssNMR experiments but without sulfoacetyl DOTOPA. A 10 μl aliquot of the solution was deposited on a circular sapphire plate (0.5 mm thickness, 25 mm diameter) and covered with an identical sapphire plate. Phase-separated droplets in the solution were imaged at room temperature using an optical microscope (Nikon SMZ-2B) with illumination from below. The sample contained between sapphire plates was then plunged into liquid nitrogen, freezing to 77 K in approximately 10 s, and stored in a liquid nitrogen bath. Imaging at low temperatures was carried out in a nitrogen-filled glove box, with the sample resting on a copper plate (10 cm × 15 cm × 6.4 mm) that had been pre-cooled to 77 K in liquid nitrogen. A 1.3 cm diameter hole in the copper plate allowed light to pass through.

### Simulations of ssNMR crosspeaks

Trajectories from molecular dynamics (MD) simulations of residues 1-163 of FUS (FUS_1-163_) in explicit water with 150 mM NaCl at 300 K were kindly provided by Drs. S. Mukherjee and L.V. Schäfer (12). From a 100 ns simulation at 25 mg/mL FUS_1-163_ concentration, atomic coordinates of one FUS_1-163_ molecule were extracted at 0.315 ns intervals (317 molecular structures), using VMD software (75). Sparta+ software (76) was then used to calculate backbone and sidechain torsion angles (ϕ, ψ, and χ_1_) and to predict ^15^N and ^13^C chemical shifts for each structure. Custom programs were used to convert the Sparta+ output files into tables that contained the torsion angles and predicted chemical shifts for all Ser, Thr, Gln, Tyr, and Gly residues in all structures, and then to generate Ramachandran surfaces and 2D ssNMR crosspeaks (*i.e.*, 2D histograms of ϕ,ψ pairs and pairs of predicted chemical shifts) for each residue type. The resulting surfaces were plotted with Origin 2024 software. Total numbers of Ser, Thr, Gln, Tyr, and Gly residues in this set of FUS_1-163_ structures were N_T_ = 12997, 3170, 11729, 7608, and 8559, respectively.

To examine the effects of changes in conformational distributions on the 2D ssNMR crosspeaks, crosspeak contributions from residues with backbone torsion angles satisfying both -180° < ϕ < -110° and 120° < ψ < 180° (encompassing β-strand-like conformations) were multiplied by a factor ε_β_. With N_β_ defined as the number of these residues for a given residue type, crosspeak contributions from all other residues of the same type were multiplied by (N_T_ - ε_β_N_β_)/ (N_T_ - N_β_) in order to maintain constant crosspeak volumes. With this approach, signal amplitudes in the difference between the 2D crosspeak with ε_β_ = 1 and the 2D crosspeak with ε_β_ > 1 are directly proportional to ε_β_ - 1. Thus, it was sufficient to calculate difference crosspeaks for a single value of ε_β_, which was chosen to be 1.2 to represent a 20% enhancement of β-strand-like conformations.

## Results

### Verification of LLPS and preservation of phase-separated droplets upon freezing

LLPS by full-length FUS, FUS-LC, and related constructs has been investigated previously by several groups (7–11,28,36,39,54). Temperatures and concentrations at which LLPS occurs are known to be sensitive to the solvent composition (43,77,78). It is therefore important to demonstrate that LLPS occurs in the glycerol-containing solvent used for ssNMR measurements and that the phase-separated state is preserved in frozen solutions.

Figs. 1c-1e show results from measurements of the temperature-dependent turbidities of FUS-LC and FUS_61-214_ solutions with compositions that match those of ssNMR samples. Measurements were performed with the device described above and depicted in Fig. S1. Phase separation, indicated by reductions in light transmission through the solutions, occurred reversibly at temperatures in the 20-40° C range. T_LLPS_ was defined as the inflection point between nearly constant light transmission above T_LLPS_ and transmission that decreases monotonically with decreasing temperature below T_LLPS_. Apparent T_LLPS_ values upon heating were generally higher by 1-3° C than T_LLPS_ values upon cooling. We attribute this hysteresis to the time required for the protein concentration to become uniform within a sample well as droplets dissolve during the heating period, especially after droplets settle towards the bottom of the sample well during the cooling period.

Interestingly, deuteration of the solvent has a significant effect on T_LLPS_. For 5.0 mg/mL FUS-LC, T_LLPS_ = 23.5 ± 1.0° C in a buffer with a 20:80 volume ratio of deuterated glycerol and H_2_O (Fig. 1c), while T_LLPS_ = 35.5 ± 1.0° C in a buffer with a 20:70:10 volume ratio of deuterated glycerol, H_2_O, and D_2_O (Fig. 1d). We attribute the higher T_LLPS_ value in the more highly deuterated solvent to an enhancement of protein-protein hydrophobic interactions, which favor phase separation, as hydrophobic interactions have been shown to be stronger in D_2_O than in H_2_O (79,80). For 14 mg/mL FUS_61-214_, T_LLPS_ = 22.5 ± 1.0° C in a buffer with a 20:70:10 volume ratio of deuterated glycerol, H_2_O, and D_2_O (Fig. 1e).

To verify that freezing does not disrupt phase-separated FUS-LC droplets, phase-separated solutions were prepared and imaged at room temperature, then frozen by immersion in liquid nitrogen and imaged while frozen. Optical imaging in the frozen state was enabled by the cryogenic imaging platform described above and depicted in Fig. S5. The resulting microscope images in Figs. 1f and 1g show droplets with diameters that range from below 5 μm to approximately 30 μm. The distribution of droplet diameters is similar at room temperature and after freezing.

Immersion in liquid nitrogen freezes the solution in a few seconds, with a cooling rate that is at least 10^4^ times slower than rates achieved with the rapid freezing apparatus used to prepare frozen solutions for ssNMR measurements. Since phase-separated droplets are preserved in a relatively slow freezing process, as shown in Figs. 1f and 1g, we conclude that the droplets must also be preserved when freezing is much more rapid. In this context, it should be noted that the translational diffusion constant for monomeric FUS-LC in 20% glycerol at 24° C is on the order of 30 μm^2^/s (81). Diffusion over distances greater than 10 μm, which would be required for dissolution of preformed droplets, would require times longer than 1 s at 24° C, even if the phase-separated state suddenly became unstable. Since the solvent viscosity increases rapidly with decreasing temperature (82), dissolution of droplets would be prohibitively slow below 0° C.

### ssNMR measurements on rapidly frozen U-^15^N,^13^C-FUS-LC solutions

Fig. 2 shows DNP-enhanced 2D ^13^C-^13^C ssNMR spectra of frozen solutions of U-^15^N,^13^C-FUS-LC at 25 K. Solutions containing 5.0 mg/ml U-^15^N,^13^C-FUS-LC and 20% glycerol by volume were rapidly frozen from temperatures well below (4° C, Fig. 2a) or well above (50° C, Fig. 2b) T_LLPS_. Although the 2D spectra do not have sufficient resolution to resolve signals from individual residues, signals arising from certain prevalent and spectroscopically distinct amino acid types can be identified. These signals, principally arising from Gly, Gln, Thr, Ser, and Tyr residues, are labeled in Figs. 2a and 2b. Chemical shift values for these amino acids (Table S2) are consistent with the values predicted for a random coil (83), indicating that FUS-LC remains largely unstructured both in the single-phase, high temperature solution and in the phase-separated, low temperature state.

**Figure 2:**
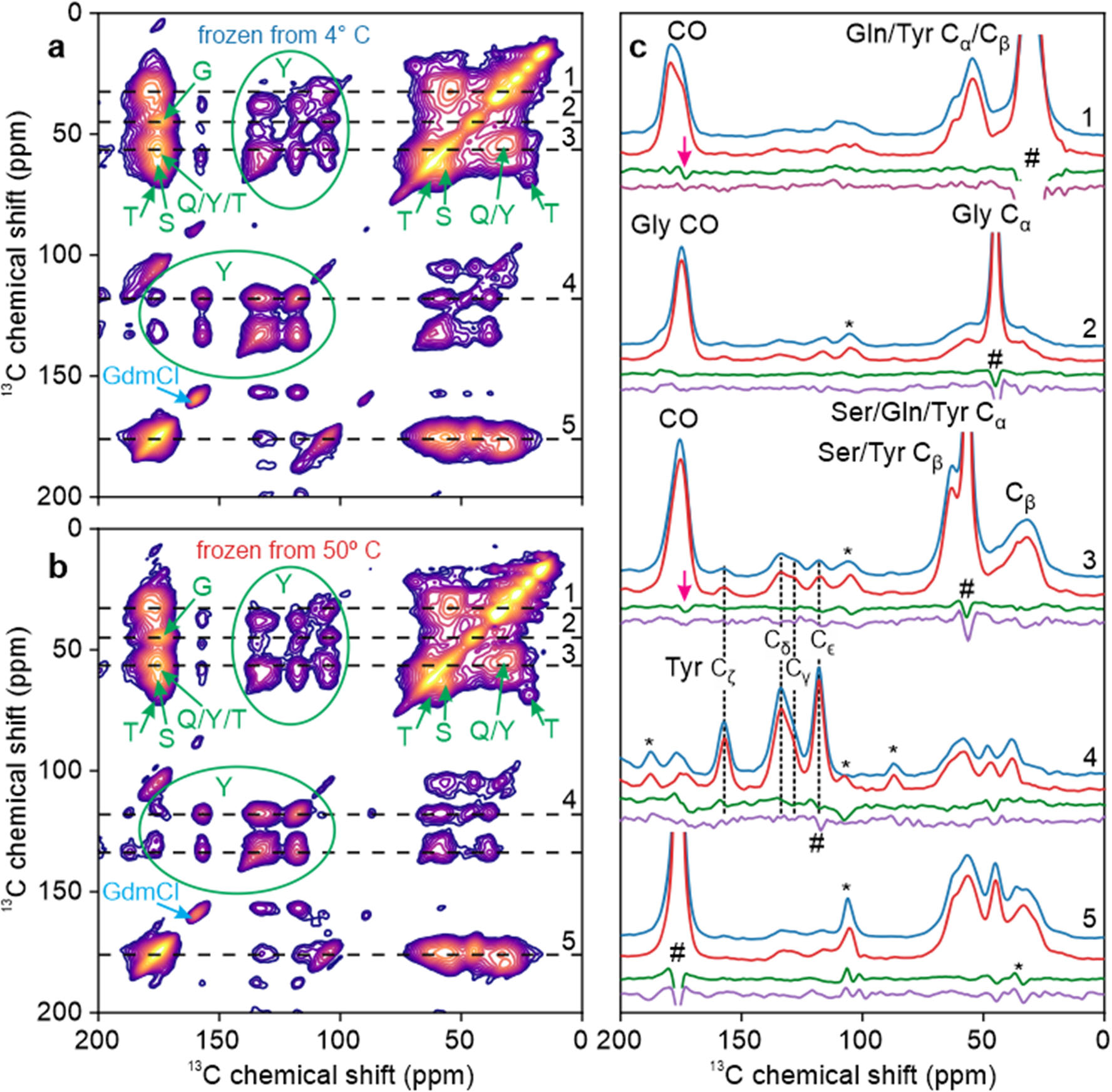
Comparison of DNP-enhanced 2D ^13^C-^13^C ssNMR spectra of U-^15^N,^13^C-FUS-LC solutions. Solutions were rapidly frozen from temperatures well below (4° C, panel a) or well above (50° C, panel b) the phase-separation temperature T_LLPS_. Signals from Gly, Gln, Thr, Tyr, and Ser residues are labeled in green. Signal from residual GdmCl is labeled in blue. (c) 1D slices from the 2D spectra in panels a and b (blue and red lines, respectively). Difference spectra for each pair of horizontal slices are shown in green, along with difference spectra for the corresponding vertical slices in purple. Signals arising from diagonal peaks and from spinning sidebands are indicated with # and * symbols, respectively. Magenta arrows indicate features that are above the noise levels in both difference spectra.

1D slices through crosspeaks in the 2D spectra are shown in Fig. 2c. To account for differences in sample packing and DNP enhancements, vertical scales of 1D slices from Fig. 2b (shown in red) were adjusted to minimize the RMS differences from signals in corresponding 1D slices from Fig. 2a (shown in blue). Comparisons of the 1D slices show that crosspeak positions and lineshapes are nearly identical in the two 2D spectra. To highlight minor differences between the spectra, 1D difference spectra were calculated for each pair of slices, both for horizontal slices (direct dimensions of the 2D spectra, difference spectra shown in green) and for the corresponding vertical slices (indirect dimensions, difference spectra shown in purple). Difference spectra are slices from Fig. 2a minus slices from Fig. 2b, after optimal scaling of slices from Fig. 2b.

Certain signals that exceed the noise levels in the 1D difference slices arise from strong diagonal signals in the 2D spectra (pound symbols in Fig. 2c). We consider these differences to be insignificant because diagonal signals were excluded from calculations of optimal scale factors. Other signals above the noise levels arise from MAS sidebands (asterisks in Fig. 2c). A small number of signals, indicated by arrows in Fig. 2c, appear to represent real differences between the 2D spectra of solutions that were frozen from temperatures above and below the phase-separation temperature, especially because these signals are present in difference spectra calculated from both horizontal and vertical slices. These signals may indicate real differences in conformational distributions at some residues of FUS-LC. In any case, these differences constitute only 3-5% of the relevant peak areas in the 1D slices (estimated by integrating the absolute values of the difference spectra) and therefore do not indicate major changes in conformational distributions (see Discussion below).

Values of DNP enhancement factors ε_DNP_ and build-up times T_DNP_ were significantly different for the samples used to obtain 2D spectra in Figs. 2a and 2b, although the two samples differed only in the incubation temperature from which they were rapidly frozen (Table S1). This observation, together with the optical microscope images in Figs. 1f and 1g, supports the idea that our rapid freezing method preserves the distinction between homogeneous and phase-separated FUS-LC solutions.

### ssNMR measurements on rapidly frozen TYG-^15^N,^13^C-FUS-LC solutions

Spectra in Fig. 2 show significant overlap between peaks from different residue types, as expected in 2D spectra of a uniformly ^13^C-labeled protein in frozen solution. In order to reduce spectral crowding, TYG-^15^N,^13^C-FUS-LC was prepared by expression in a defined amino acid medium containing ^15^N,^13^C-labeled Thr and Tyr residues. All other amino acids were provided at natural isotopic abundance, but Gly residues in the resulting protein sample were partially labeled through scrambling from labeled Thr or Tyr. Fig. 3 shows DNP-enhanced 2D ^13^C-^13^C ssNMR spectra at 25 K of TYG-^15^N,^13^C-FUS-LC solutions that were rapidly frozen after incubation at temperatures well below (4° C, Fig. 3a) or well above (50° C, Fig. 3b) T_LLPS_. Crosspeaks from Thr, Tyr, and Gly residues are resolved in these 2D spectra, aside from overlap of the CO/C_α_ crosspeaks from Thr and Tyr residues. Full-width-at-half-maximum (FWHM) linewidths are 6.5-8.0 ppm, significantly greater than the 2-4 ppm linewidths that are typically observed for individual residues in ^13^C ssNMR spectra of frozen solutions of folded proteins under the same measurement conditions (67,68,84,85). We attribute the linewidths in Fig. 3 to a combination of two factors: (i) FUS-LC adopts a broad distribution of conformations, which do not exchange among themselves in frozen solutions. Therefore, the full conformational dependences of ^13^C chemical shifts (86–88) are preserved as inhomogeneous broadening in these spectra (89,90); (ii) Our FUS-LC sequence (including the N-terminal tag) contains 12 Thr, 31 Tyr, and 52 Gly residues, all of which contribute to the crosspeak signals from each residue type.

**Figure 3:**
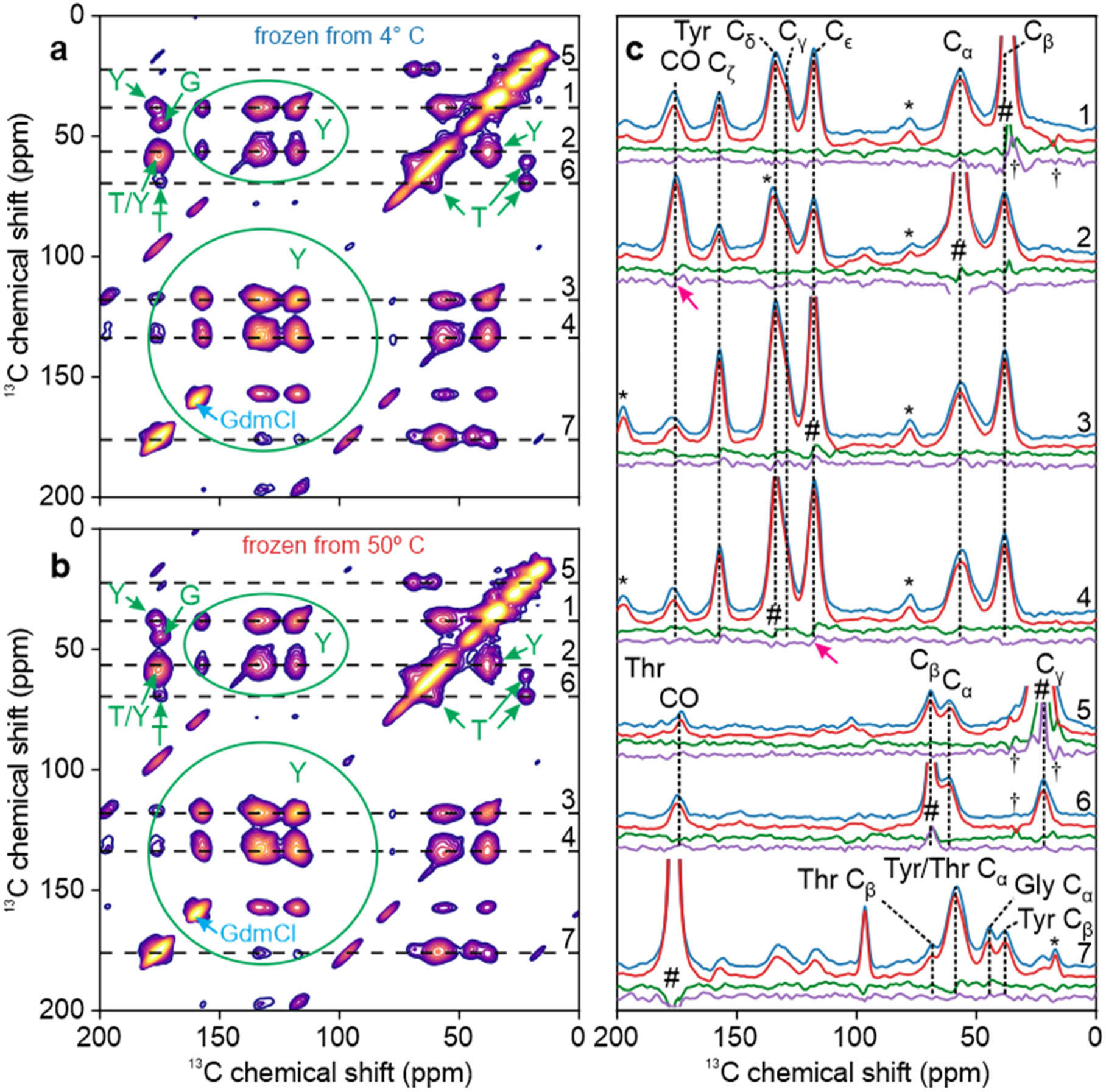
Comparison of DNP-enhanced 2D ^13^C-^13^C ssNMR spectra of TYG-^15^N,^13^C-FUS-LC solutions. Solutions were rapidly frozen from temperatures well below (4° C, panel a) or well above (50° C, panel b) T_LLPS_. Signals from Thr, Tyr, and Gly residues are labeled in green. Signal from residual GdmCl is labeled in blue. (c) 1D slices from the 2D spectra in panels a and b (blue and red lines, respectively). Difference spectra for each pair of horizontal slices are shown in green, along with difference spectra for the corresponding vertical slices in purple. Signals arising from diagonal peaks and from spinning sidebands are indicated with # and * symbols, respectively. Artifacts caused by t_1_ noise from large natural-abundance ^13^C signals of residual frozen isopentane are indicated with † symbols. Magenta arrows indicate features that are above the noise levels in both difference spectra.

Fig. 3c shows 1D slices through crosspeaks in the 2D spectra in Figs. 3a and 3b, along with 1D difference spectra for horizontal and vertical slices, as in Fig. 2c. Arrows indicate features of the difference spectra that are above the noise level in both the horizontal and vertical directions. These features are small, constituting approximately 5-8% of the relevant peak areas in the 1D slices. Thus, major differences between conformational distributions of Thr residues, Tyr residues, and Gly residues of FUS-LC in homogeneous and phase-separated solutions are not detected even when overlap between signals from different residue types is minimized.

2D ^15^N-^13^C spectra of the same rapidly frozen TYG-^15^N,^13^C-FUS-LC solutions are shown in Figs. 4a and 4b. 1D slices parallel to the ^13^C and ^15^N chemical shift axes, as well as the corresponding 1D difference spectra, are shown in Figs. 4c and 4d, respectively. No signals are observed that are significantly above the noise levels in the difference spectra.

**Figure 4:**
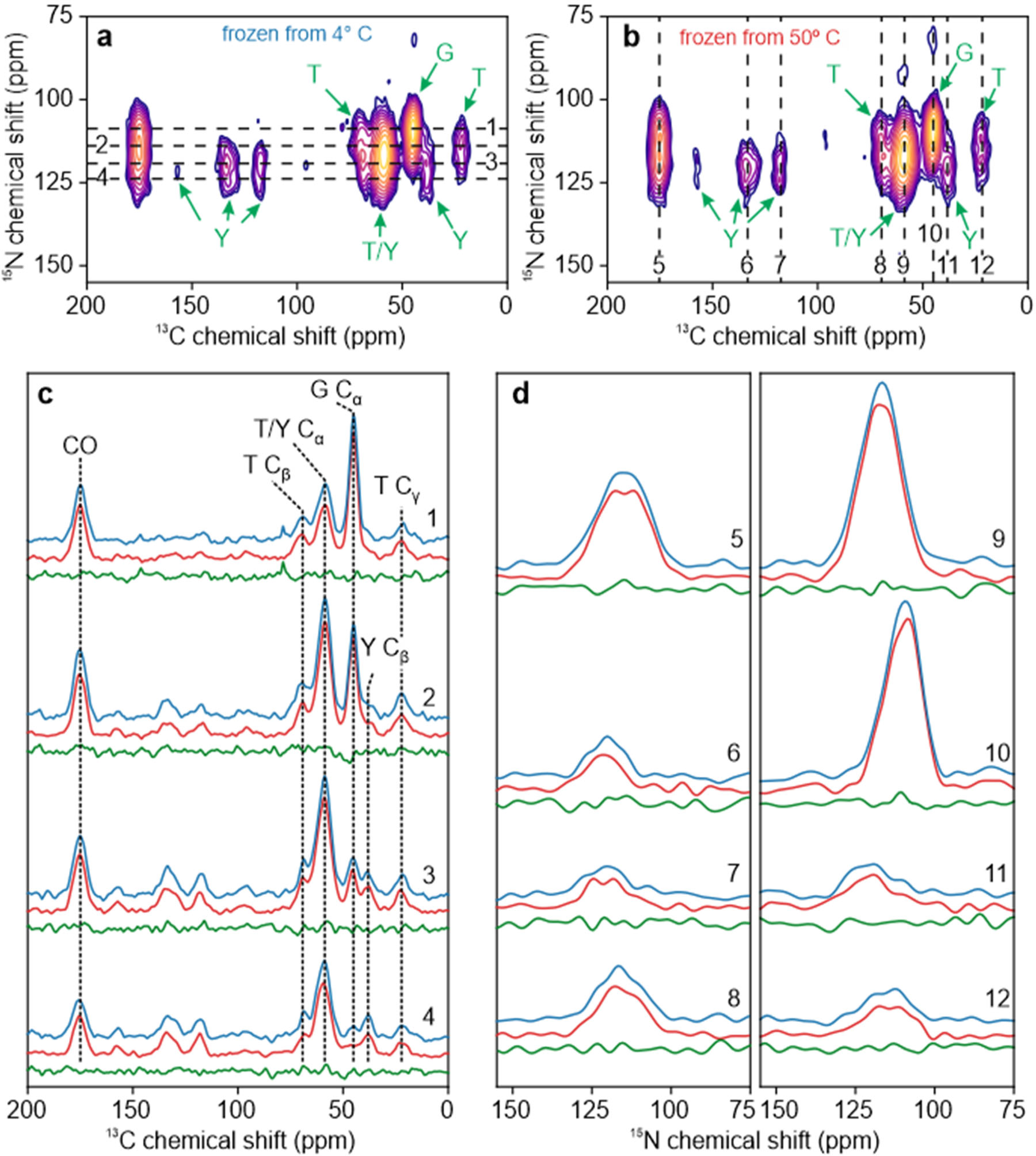
Comparison of DNP-enhanced 2D ^15^N-^13^C ssNMR spectra of TYG-^15^N,^13^C-FUS-LC solutions. Solutions were rapidly frozen from temperatures well below (4° C, panel a) or well above (50° C, panel b) T_LLPS_. Signals from Thr, Tyr, and Gly residues are labeled in green. (c) Horizontal 1D slices from the 2D spectra in panels a and b (blue and red lines, respectively). Slices were taken at the positions shown in panel a. Difference spectra for each pair of horizontal slices are shown in green. (d) Vertical 1D slices from the 2D spectra in panels a and b (blue and red lines, respectively). Slices were taken at the positions shown in panel b. Difference spectra for each pair of vertical slices are shown in green.

### ssNMR measurements on rapidly frozen sel-^15^N,^13^C-FUS_61-214_ solutions

One path to resolution of ssNMR signals from individual residues is selective isotope labeling, which can be achieved through native chemical ligation (63,64). To this end, we prepared sel-^15^N,^13^C-FUS_61-214_ by ligating FUS_61-92_, chemically synthesized with ^15^N,^13^C-labeling at G65, Q69, T78, Y81, and S86, to recombinantly expressed FUS_93-214_, with the Gln-to-Cys substitution at residue 93 required for ligation. As shown in Fig. 1e, this construct exhibits LLPS at 22.5 ± 1.0° C at the concentration and with the solvent used for ssNMR measurements.

Fig. 5 shows DNP-enhanced 2D ^13^C-^13^C ssNMR spectra at 25 K of sel-^15^N,^13^C-FUS_61-214_ solutions that were rapidly frozen after incubation at temperatures well below (4° C, Fig. 5a) or well above (50° C, Fig. 5b) T_LLPS_. Although CO/C_α_ crosspeaks from Q69, T78, Y81, and S86 overlap in these 2D spectra, other crosspeaks are resolved and represent signals from individual sites in the ligated construct. ^13^C chemical shift values for the five labeled residues in sel-^15^N,^13^C-FUS_61-214_ (Table S3) are similar to the values measured for the identifiable amino acids in U-^15^N,^13^C-FUS-LC (Table S2), indicating that the labeled residues in FUS_61-214_ adopt conformational distributions that are similar to those in full-length FUS-LC.

**Figure 5:**
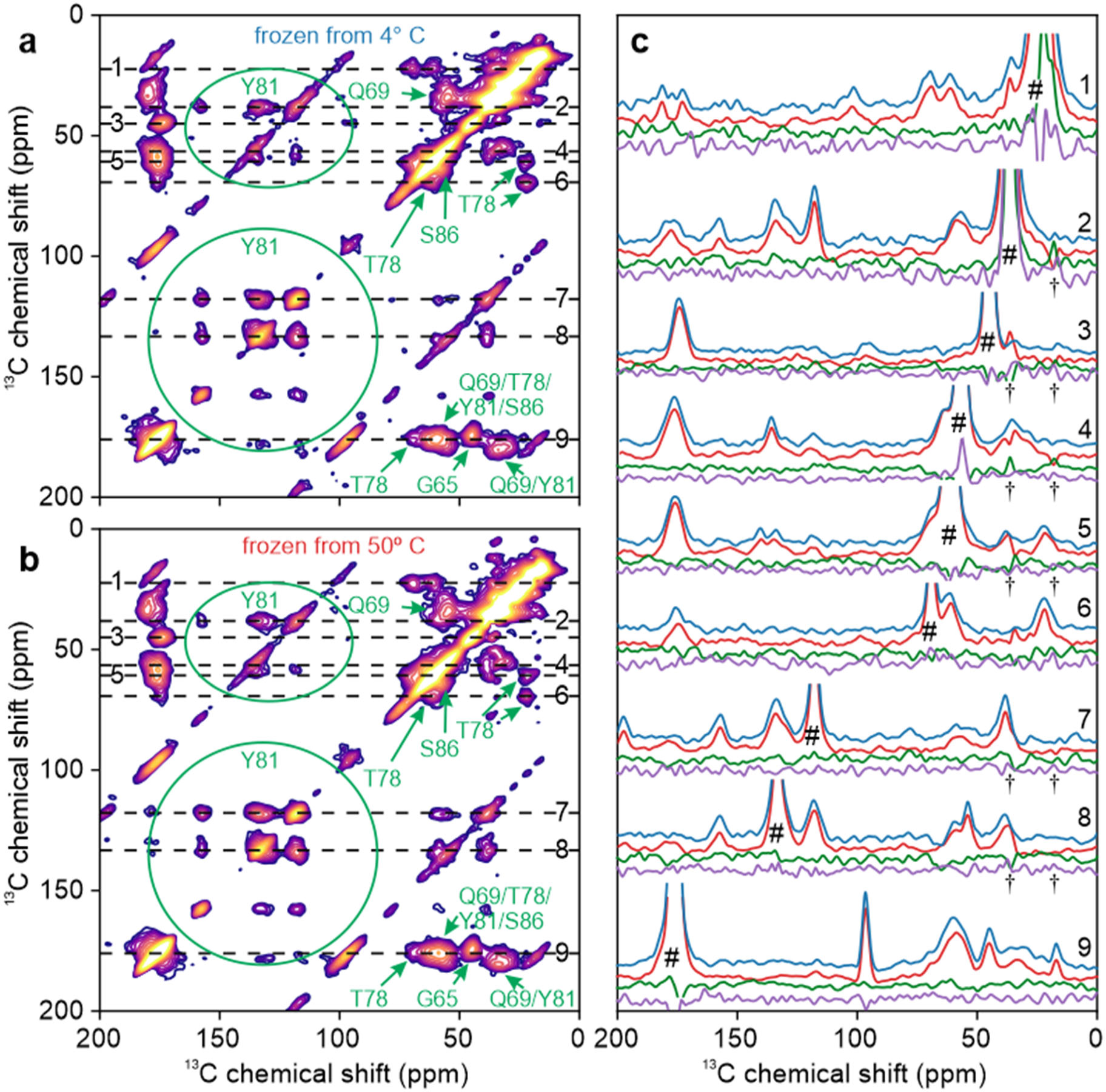
Comparison of DNP-enhanced 2D ^13^C-^13^C ssNMR spectra of sel-^15^N,^13^C-FUS_61-214_ solutions. Solutions were rapidly frozen from temperatures well below (4° C, panel a) or well above (50° C, panel b) T_LLPS_. Signals from G65, Q69, T78, Y81, and S86 are labeled in green. (c) 1D slices from the 2D spectra in panels a and b (blue and red lines, respectively). Difference spectra for each pair of horizontal slices are shown in green, along with difference spectra for the corresponding vertical slices in purple. Signals arising from diagonal peaks are indicated with # symbols. Artifacts caused by t_1_ noise from large natural-abundance ^13^C signals of residual frozen isopentane are indicated with † symbols.

Fig. 5c shows 1D slices through crosspeaks in the 2D spectra, along with difference spectra extracted along the direct and indirect dimensions. No features above the noise level are observed in the 1D difference spectra, other than features due to small differences in diagonal signals or t_1_ noise from natural-abundance ^13^C signals of frozen isopentane that was not completely removed during sample packing.

Signal-to-noise ratios (SNRs) in 2D ^13^C-^13^C ssNMR spectra of frozen sel-^15^N,^13^C-FUS_61-214_ solutions are lower than SNRs in 2D spectra of frozen U-^15^N,^13^C-FUS-LC and TYG-^15^N,^13^C-FUS-LC solutions for two reasons. First, the number of isotopically labeled residues is much smaller. Second, ssNMR measurements on frozen sel-^15^N,^13^C-FUS_61-214_ solutions used a 200 mW microwave source (rather than 1.5 W source) for DNP (see Materials and Methods), resulting in lower signal enhancements from DNP. Motivated by the lower SNRs of 2D spectra in Figs. 5a and 5b, we sought to identify small differences between these 2D spectra by comparing the centers-of-mass of well-resolved crosspeaks. Center-of-mass coordinates were calculated (see Materials and Methods) for spectral intensities within the regions indicated in Fig. 6a by solid-line rectangles (above the diagonal) and for the symmetric regions indicated by dashed-line rectangles (below the diagonal). The results are plotted in Fig. 6b, with blue symbols for centers-of-mass from Fig. 5a and red symbols for centers-of-mass from Fig. 5b. For Fig. 6b, coordinates in the direct and indirect dimensions are swapped for crosspeaks below the diagonal, and the coordinates are plotted relative to the average values for each crosspeak.

**Figure 6:**
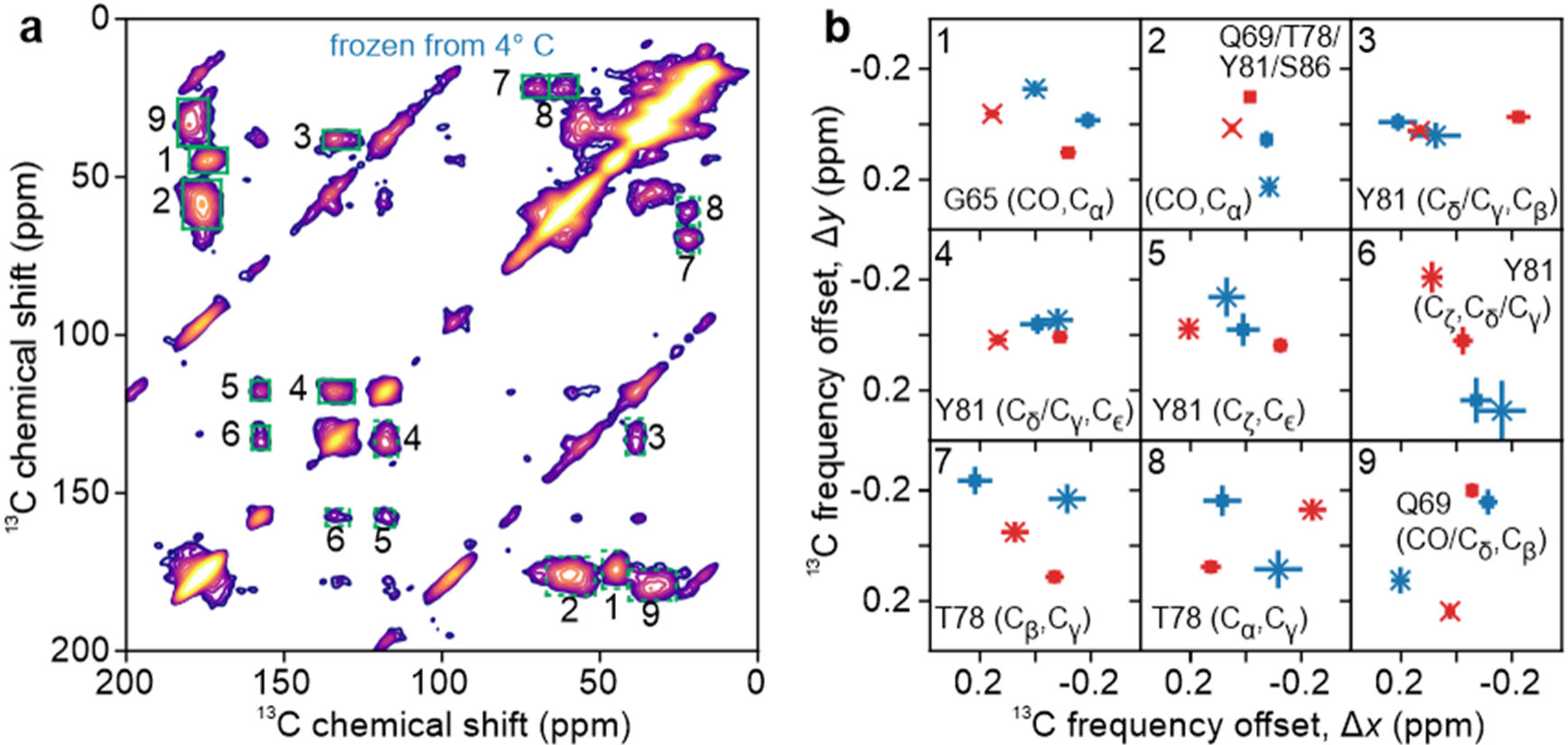
Center-of-mass analysis of 2D spectra. (a) DNP-enhanced 2D ^13^C-^13^C ssNMR spectrum of a sel-^15^N,^13^C-FUS_61-214_ solution that was rapidly frozen 4° C (same as Fig. 5a). Center-of-mass values were calculated for crosspeak signals within green rectangles. Solid-line and dashed-line rectangles with the same number are related by symmetry across the diagonal of the 2D spectrum. (b) Center-of-mass values of the indicated crosspeaks, calculated for 2D spectra in Fig. 5a (blue) and Fig. 5b (red) and plotted relative to the average values. Values calculated for rectangular regions above and below the diagonal are shown with ▪ and × symbols, respectively. Bars indicate uncertainties, calculated as explained in the text.

Center-of-mass coordinates for symmetric pairs of crosspeaks above and below the diagonal (squares and crosses, respectively, in Fig. 6b) would ideally be identical, but in fact differ by as much as 0.4 ppm. In contrast, uncertainties estimated purely from the SNR values of the 2D spectra, represented by bars in Fig. 6b, are 0.2 ppm or less. Thus, uncertainties in center-of-mass coordinates are apparently determined largely by baseline imperfections and spurious signal contributions from t_1_ noise. Moreover, differences between the calculated center-of-mass coordinates of crosspeaks in Figs. 5a and 5b are similar in size to the differences between center-of-mass coordinates of crosspeaks above and below the diagonal in each individual spectrum. We therefore conclude that any differences in the centers-of-mass for crosspeaks in the two 2D spectra of frozen solutions of sel-^15^N,^13^C-FUS_61-214_ are within the uncertainties of our measurements.

Center-of-mass coordinates were also calculated for crosspeaks in the 2D ^13^C-^13^C ssNMR spectra of frozen solutions of TYG-^15^N,^13^C-FUS-LC shown in Figs. 3a and 3b. Again, differences between center-of-mass coordinates for crosspeaks in the two 2D spectra were similar in size to differences between center-of-mass coordinates for crosspeaks above and below the diagonal in each individual spectrum (Fig. S6). These differences are therefore within the uncertainties of our measurements.

## Discussion

Experiments described above were designed to test the hypothesis that LLPS by an IDP, specifically the LC domain of FUS, is accompanied by changes in local conformational distributions. Such changes might conceivably result from specific conformational requirements of certain intermolecular interactions that may contribute to the lower free energy of the phase-separated state of the IDP solution below T_LLPS_, such as intermolecular backbone hydrogen bonds that favor β-strand-like conformations or intermolecular hydrogen bonds between sidechains of Gln residues that favor certain sets of sidechain torsion angles. The fact that NMR chemical shifts, especially ^13^C chemical shifts and including chemical shifts in ssNMR spectra, are sensitive to local conformational details (86–88) suggests that changes in conformational distributions would have detectable effects on ssNMR spectra (89,90).

As summarized in the Introduction, the possibility that conformational changes occur in phase-separated states of IDPs has been examined in previous studies, including studies by solution NMR (8,9,11,19–21,39–49), EPR (11,52–54), computational methods (12,14,22,23,56), and biochemical approaches (28,57–59). For the most part, studies based on solution NMR methods have not found evidence for large changes in the time-averaged conformations of IDPs or for exchange between unstructured and highly structured components in the phase-separated state. Other experimental studies have provided some evidence for molecular structural changes that accompany LLPS (11,21,47,51,53,54).

Experiments on FUS-LC and FUS_61-214_ described above use an approach that has not been previously applied to the issue of conformational changes that may accompany LLPS. Noteworthy aspects of this approach include: (i) To our knowledge, these are the first experiments in which 2D crosspeak lineshapes in ssNMR spectra of frozen solutions are used to compare conformational distributions of an IDP in its homogeneous and phase-separated states. Lineshapes in these spectra are not affected by exchange among conformations (on any time scale) or by other molecular motions, since conformational exchange and molecular motions (other than small-amplitude vibrations and librations) are quenched at the 25 K sample temperatures of our measurements; (ii) Our experiments directly compare homogeneous and phase-separated solutions under conditions that are otherwise identical. Total protein concentrations, solvent compositions, and sample temperatures are the same in measurements on the homogeneous samples (Figs. 2b, 3b, 4b, and 5b) and the corresponding phase-separated samples (Figs. 2a, 3a, 4a, and 5a). Thus, differences in solvent conditions and temperatures cannot produce differences in the ssNMR spectra; (iii) All isotopically labeled sites and all components of their conformational distributions contribute to the 2D ssNMR spectra with approximately equal weighting. Minor conformation-dependent variations in signal intensities may arise from variations in ^1^H-^13^C cross-polarization efficiencies (or ^1^H-^15^N and ^15^N-^13^C cross-polarization efficiencies in the case of the 2D ^15^N-^13^C spectra in Fig. 4) or from local variations in DNP-enhanced ^1^H spin polarizations. However, the resulting conformation-dependent variations in signal intensities should not exceed 10% when molecular motions are quenched at 25 K; (iv) Before rapid freezing, the phase-separated droplets in our experiments are formed at 4° C, where the protein concentration and viscosity are higher than at temperatures closer to T_LLPS_. Changes in local conformational distributions accompanying LLPS are likely to be larger when the droplets are formed well below T_LLPS_, rather than slightly below T_LLPS_.

Experiments on U-^15^N,^13^C-FUS-LC result in nearly identical 2D ^13^C-^13^C spectra (Figs. 2a and 2b). The only significant differences (*i.e.*, differences that are above the noise levels in both horizontal and vertical 1D slices and are not attributable to artifacts) are in 1D slices through ^13^CO signals (Fig. 2c). These differences are at the 3-5% level relative to the total peak areas. 2D ssNMR spectra of TYG-^15^N,^13^C-FUS-LC exhibit less overlap between crosspeaks, especially in the aliphatic ^13^C chemical shift region, due to the limited number of isotopically labeled residues. In principle, the reduction in overlap could facilitate the detection of differences between spectra of samples that were frozen from 4° C or 50° C. In fact, the 2D spectra are still very similar (Figs. 3a and 3b, Figs. 4a and 4b), showing differences only at the 5-8% level in 1D slices (Fig. 3c). To obtain 2D spectra with resolved crosspeaks from individual amino acid sites, we prepared sel-^15^N,^13^C-FUS_61-214_ by ligating a synthetic N-terminal segment, containing five labeled residues, with an unlabeled C-terminal recombinant segment. 2D ^13^C-^13^C ssNMR spectra of sel-^15^N,^13^C-FUS_61-214_ solutions frozen from 4° C or 50° C (Fig. 5) exhibit no significant differences (although the SNR in these 2D spectra is lower than in 2D spectra of U-^15^N,^13^C-FUS-LC and TYG-^15^N,^13^C-FUS-LC). Based on these experimental results, we conclude that LLPS by the LC domain of FUS does not involve major changes in local conformational distributions.

The observation of small differences in certain 1D slices from 2D ^13^C-^13^C ssNMR of frozen U-^15^N,^13^C-FUS-LC and TYG-^15^N,^13^C-FUS-LC solutions (Figs. 2c and 3c) suggests that minor differences in local conformational distributions may exist between the phase-separated and homogeneous solutions. To estimate the maximum size of such differences, we extracted conformational distributions for Ser, Thr, Gln, and Tyr residues from all-atom MD simulations on FUS_1-163_ reported by Mukherjee and Schäfer (12). Ramachandran plots from 317 FUS_1-163_ structures in a 100 ns MD trajectory are shown in Fig. S7. We used the Sparta+ program (76) to predict ^15^N and ^13^C chemical shifts for all Ser, Thr, Gln, and Tyr residues in these structures, from which we generated the simulated ^13^C_α_/^13^C_β_, ^13^C_α_/^13^CO, and ^15^N/^13^C_α_ crosspeaks shown in Fig. 7. We then simulated “reweighted” crosspeaks in which the signal contributions from residues with β-strand-like conformations (defined by backbone torsion angle ranges -180° < ϕ < -110° and 180° > ψ > 120°) were enhanced by a factor of ε_β_ = 1.2, while contributions from residues with other conformations were uniformly reduced in order to maintain constant crosspeak volumes. Differences between the reweighted crosspeaks and the original crosspeaks are also shown in Fig. 7. Maximum values of the crosspeak differences are 5-18% for ^13^C_α_/^13^C_β_, 11-16% for ^13^C_α_/^13^CO, and 11-15% for ^15^N/^13^C_α_ crosspeaks. Magnitudes of minimum values are 8-15% for ^13^C_α_/^13^C_β_, 7-10% for ^13^C_α_/^13^CO, and 6-11% for ^15^N/^13^C_α_ crosspeaks. Since crosspeak differences in Fig. 7 are proportional to ε_β_-1, a comparison of these simulations with the experimental results for rapidly frozen FUS-LC solutions suggests that local conformation distributions for FUS-LC in phase-separated solutions may differ by roughly 5-10% from distributions in homogeneous solutions. In other words, relative populations within a certain range of conformations may be enhanced by 5-10% without producing changes in 2D ssNMR crosspeak shapes that exceed the experimentally observed changes.

**Figure 7:**
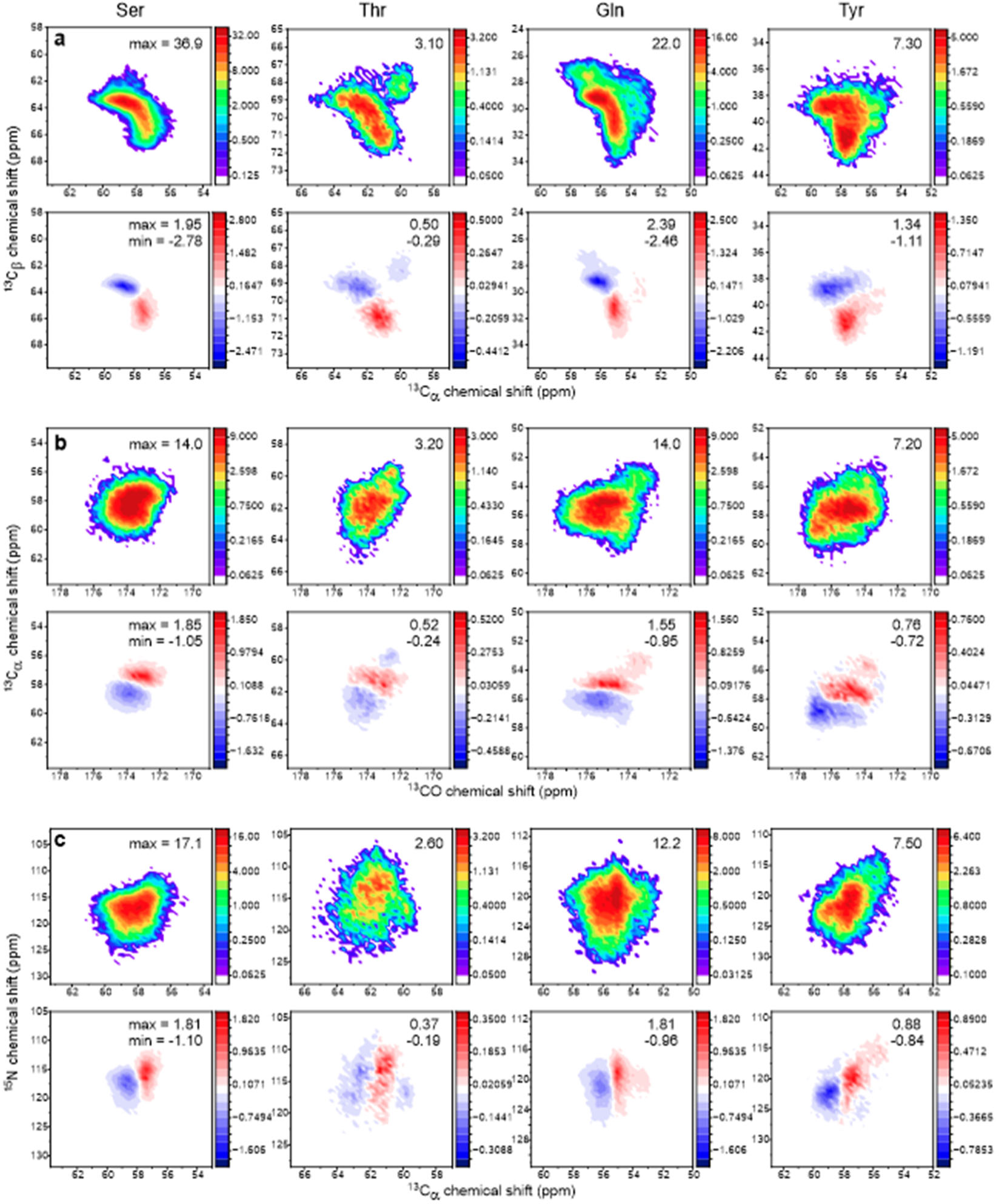
Simulated 2D ssNMR crosspeak lineshapes for Ser, Thr, Gln, and Tyr residues, based on conformational distributions from MD simulations on FUS_1-163_ (see text and Fig. S6). Crosspeaks for intraresidue ^13^C_α_/^13^C_β_, ^13^CO/^13^C_α_, and ^15^N/^13^C_α_ chemical shift correlations are shown in panels a, b, and c, respectively. Upper plots show crosspeaks calculated without reweighting of conformations. Maximum crosspeak signal intensities are indicated. Lower plots show differences between reweighted crosspeaks, in which contributions from β-strand-like conformations are enhanced by a factor of 1.2, and crosspeaks without reweighting. Maximum and minimum signal intensities in these 2D difference spectra are indicated.

Is it possible that substantial changes in local conformational distributions occur during the rapid freezing process, so that ssNMR measurements on rapidly frozen solutions do not accurately reflect the conformational properties of FUS-LC solutions in the phase-separated state near 4° C or the homogeneous state near 50° C? As shown by optical microscope images in Fig. 1, phase-separated droplets remain intact after freezing, even when freezing occurs slowly (time scale of seconds, compared with the sub-millisecond time scale for freezing of ssNMR samples). In addition, as noted above, translational diffusion of FUS-LC molecules is much too slow to permit dissolution of phase-separated droplets during rapid freezing of samples that were equilibrated at 4° C.

For samples that were equilibrated at 50° C before rapid freezing, one might ask whether FUS-LC molecules undergo LLPS during the freezing process. Time-resolved x-ray scattering experiments by Martin *et al*. (77) have shown that the time scale for phase separation of solutions of a 159-residue IDP, representing the LC domain of hnRNPA1, is greater than 10 ms when LLPS is initiated by a rapid increase in salt concentration. The x-ray scattering data show that the radius of gyration of individual protein chains decreases before oligomerization and LLPS, but the reported time scale for contraction of protein monomers is 2 ms. Given the general similarity of LC domains of hnRNPA1 and FUS, the greater viscosity of our glycerol/water solutions, the increase in viscosity as temperature decreases, and the high cooling rates in our experiments (greater than 10^5^ °/s (68)), phase separation and substantial conformational changes during rapid freezing of our FUS-LC solutions from 50° C can be safely ruled out. In addition, as discussed above, measurements of DNP enhancement factors and build-up times support a qualitative difference between rapidly frozen FUS-LC solutions that were first equilibrated at either 4° C or 50° C, even though the 2D ssNMR spectra are quite similar. It should also be noted that previous ssNMR studies in our laboratory provide evidence that rapid freezing of protein solutions under our conditions does not perturb protein structures, as no discrepancies between ssNMR data on the frozen solutions and the expected structural properties prior to freezing have been observed (67,68,84,85).

In conclusion, the ssNMR data for rapidly frozen FUS-LC and FUS_61-214_ solutions presented above indicate that local conformational distributions in these IDPs are nearly the same in homogeneous and phase-separated states. Minor differences in 2D ssNMR spectra may reflect small changes in the relative populations of certain conformations, but these changes can be no more than 5-10%. Thus, although a variety of intermolecular interactions are present in phase-separated droplets, these interactions do not require a large increase in populations of β-strand-like conformations or other specific secondary structures.

## Data and code availability

2D ssNMR data, data used to generate plots in Figs. 7 and S7, and the Fortran95 program used to generate these plots are available at https://doi.org/10.17632/ywssdrmzvp.1. All other data are available upon request from the authors at.

## Supporting material

Supporting material can be found online at XXXXX.

## Author contributions

CBW and RT designed research and analyzed data. CBW, ML, and WMY prepared samples and performed research. CBW, ML, WMY, and RT wrote the manuscript.

## Acknowledgements

This work was supported by the Intramural Research Program of the National Institute of Diabetes and Digestive and Kidney Diseases, National Institutes of Health.

