## Supplemental Material for "Conformations of a Low-Complexity Protein in Homogeneous and Phase-Separated Frozen Solutions"

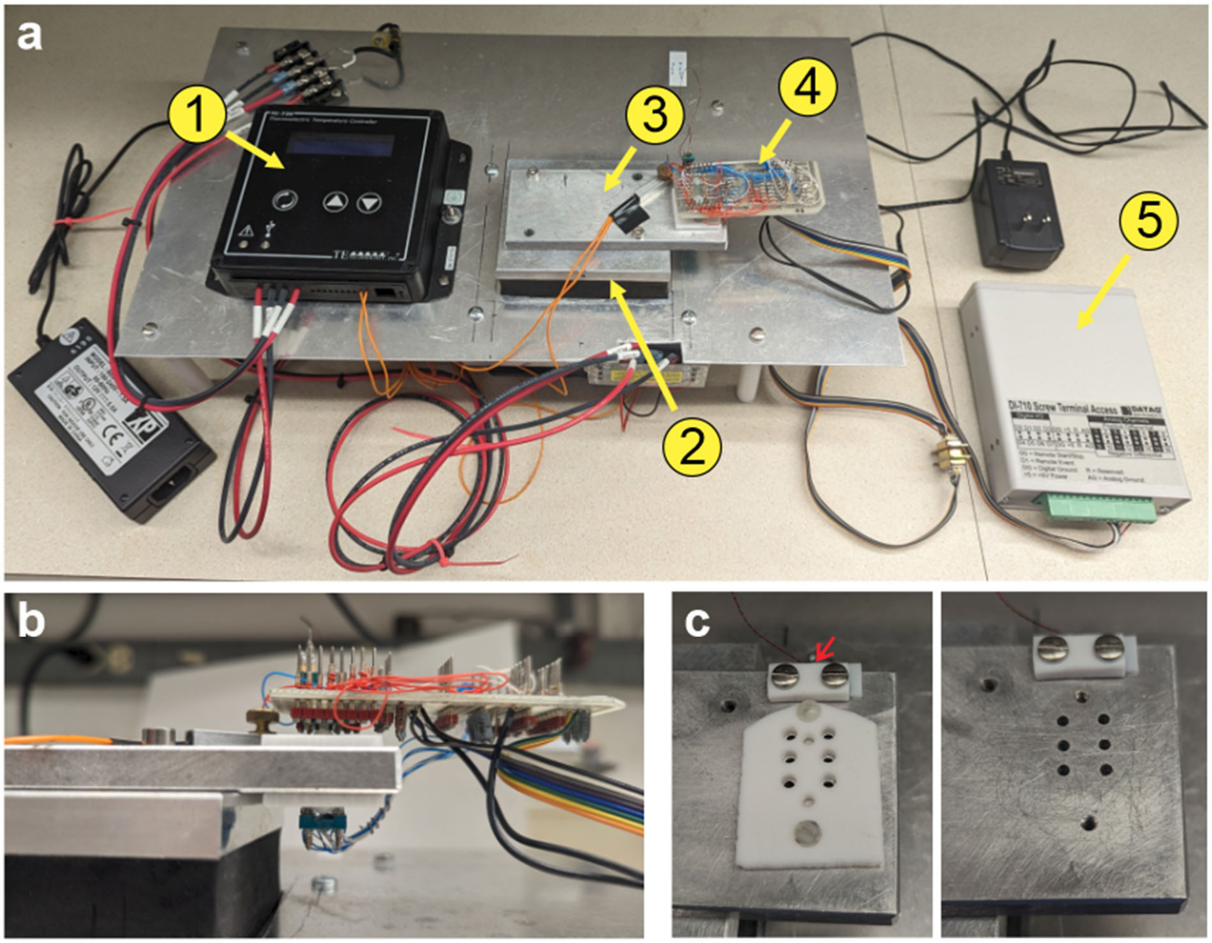


Figure S1: Apparatus for measurements of temperature-dependent turbidities, as in Fig. 1c-e. (a) Main components, consisting of a TC-720 temperature controller (1) and CP-063HT thermoelectric cooler (2) from TE Technology, Inc., an aluminum plate with six sample wells attached to the thermoelectric cooler (3), circuitry that senses the plate temperature and the transmission of 570 nm wavelength light through each sample well (4), and a DATAQ Instruments DI-710-UH analog-to-digital converter (5) that sends the temperature and transmitted light intensity data to a computer for storage and analysis. (b) Side view of the circuitry, with light-emitting diodes directly below and phototransistors directly above each sample well. (c) Views of the sample wells with (left) and without (right) a polytetrafluoroethylene (PTFE) piece that supports the detection circuitry. Red arrow points to the location of a 100 Ω Pt resistor, which is embedded in the aluminum plate and used to monitor sample temperatures.

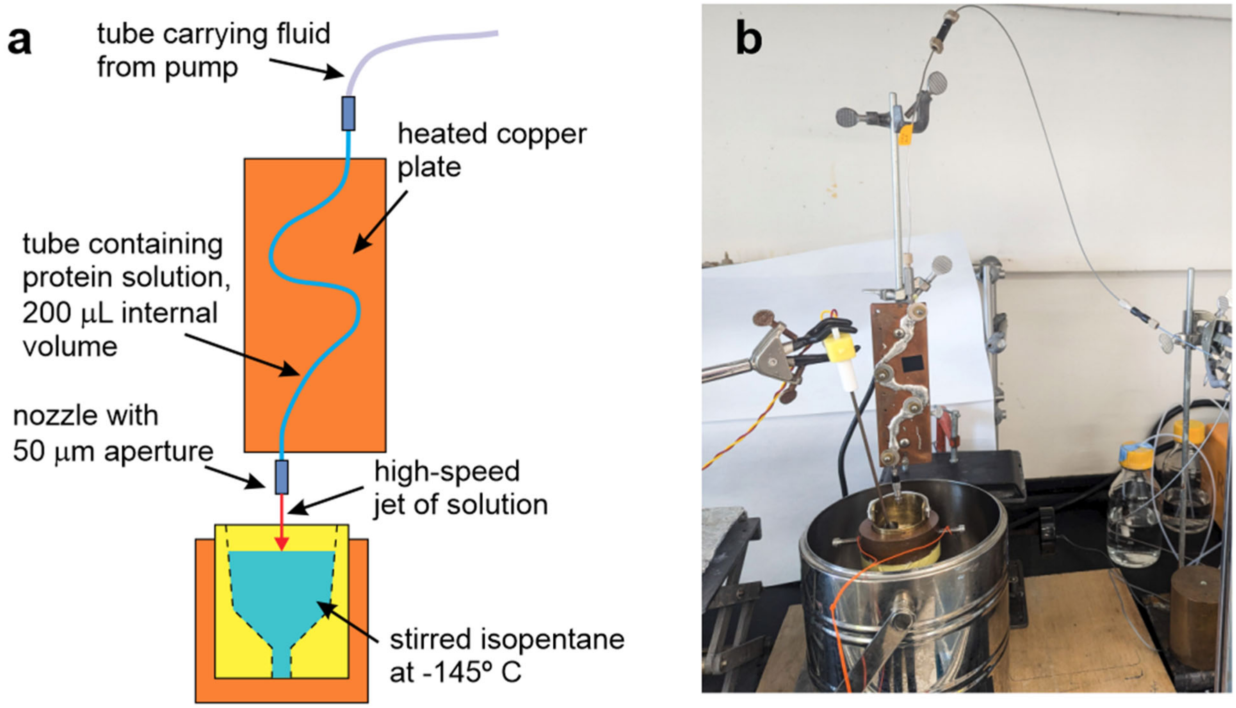


Figure S2: (a) Schematic representation of the apparatus used for rapid freezing of FUS-LC and $FUS_{61-214}$ solutions from controlled initial temperatures. The solution is first loaded into a tube that is soldered to a copper plate, which can be heated to a specified temperature. An aliquot of hexane is loaded behind the protein solution to separate the protein solution from driving fluid that is delivered by a pump. When pump pressure is applied, the protein solution is expelled through a nozzle as a high-speed jet and freezes in a stirred bath of cold isopentane. (b) Photograph of the apparatus.

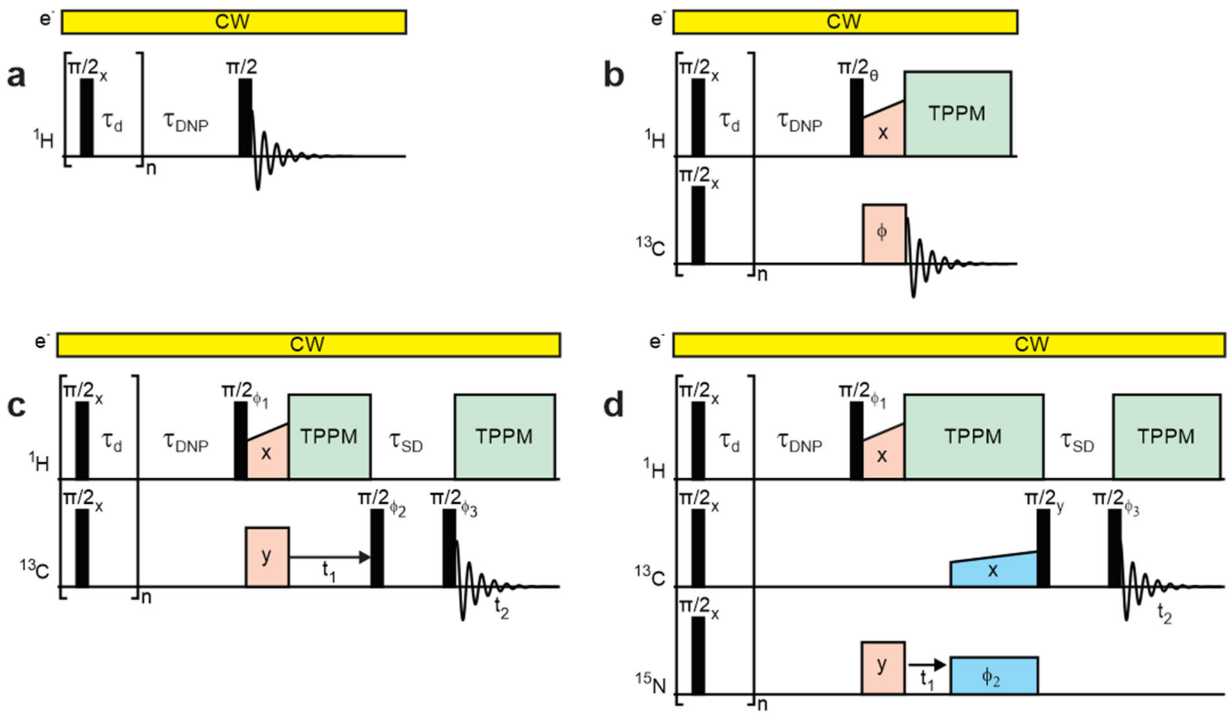


Figure S3: Pulse sequences for ssNMR measurements. Continuous-wave (CW) microwaves at 263.9 GHz are applied to electron spins (e-) to drive cross-effect DNP. Radio-frequency pulses are applied to $^{1}$H, $^{13}$C, and $^{15}$N nuclear spins at 400.8 MHz, 100.8 MHz, and 40.6 MHz, respectively. Pulse sequences begin with trains of $\pi/2$ pulses, separated by delays $\tau_D$ = 10 ms, to destroy pre-existing nuclear spin polarizations. $^{1}$H spin polarization then builds up during $\tau_{DNP}$. (a) Pulse sequence for measurement of 1D $^{1}$H ssNMR spectra. (b) Pulse sequence for measurement of 1D $^{13}$C ssNMR spectra after $^{1}$H-$^{13}$C cross-polarization (pink blocks). $^{1}$H decoupling with two-pulse phase modulation (TPPM, green blocks)) is applied during acquisition of $^{13}$C signals. Phase cycling: $\theta = y\bar{y}$; $\phi = xx\bar{x}\bar{x}yy\bar{y}\bar{y}$; receiver phase $=x\bar{x}\bar{x}xy\bar{y}\bar{y}y$. (c) Pulse sequence for measurement of 2D $^{13}$C-$^{13}$C ssNMR spectra, with $^{13}$C-$^{13}$C polarization transfers between $t_1$ and $t_2$ periods by spin diffusion during $\tau_{SD}$. Phase cycling: $\phi_1 = yy\bar{y}\bar{y}$; $\phi_2 = x\bar{x}$ or $y\bar{y}$; $\phi_3 = xxxx\bar{x}\bar{x}\bar{x}\bar{x}$; receiver phase $=x\bar{x}\bar{x}x\bar{x}xx\bar{x}$. (d) Pulse sequence for measurement of 2D $^{15}$N-$^{13}$C ssNMR spectra, with $^{1}$H-$^{15}$N cross-polarization (pink blocks) before and $^{15}$N-$^{13}$C cross-polarization (blue blocks) after the $t_1$ period. Phase cycling: $\phi_1 = y\bar{y}$; $\phi_2 = yy\bar{y}\bar{y}$ or $xx\bar{x}\bar{x}$; $\phi_3 = xxxxyyyy\bar{x}\bar{x}\bar{x}\bar{x}\bar{y}\bar{y}\bar{y}\bar{y}$; receiver phase = $x\bar{x}\bar{x}xy\bar{y}\bar{y}y\bar{x}xx\bar{x}\bar{y}yy\bar{y}$.

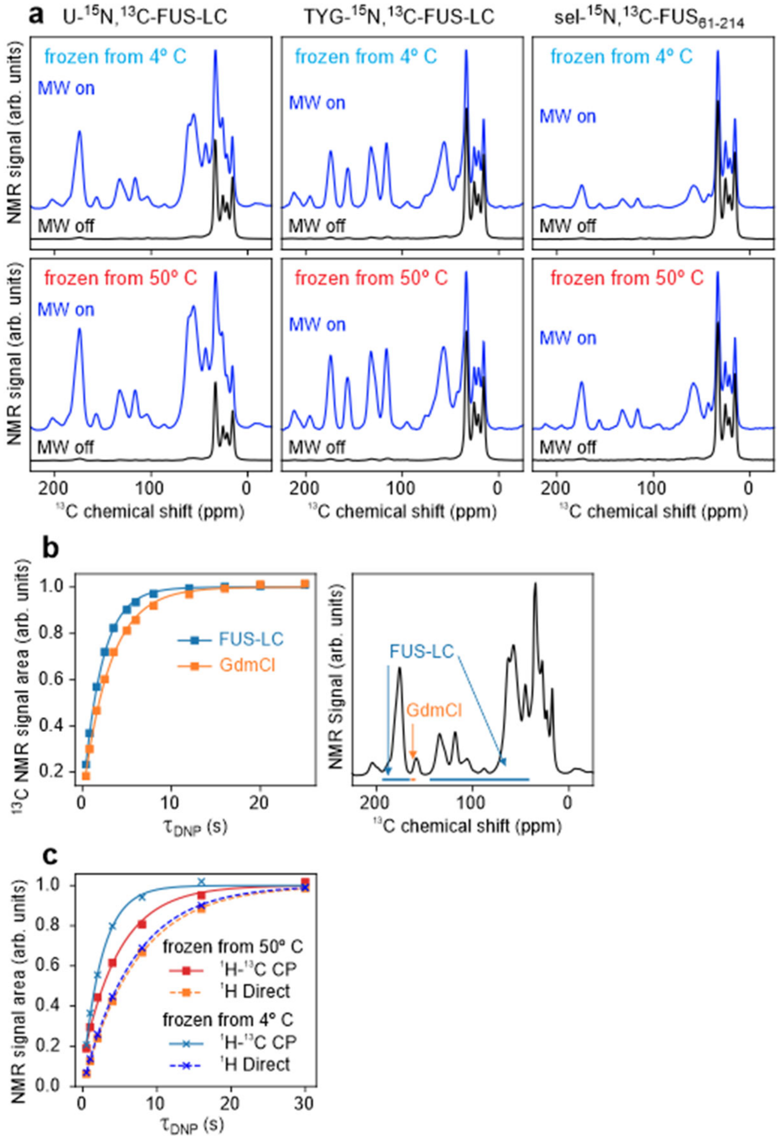


Figure S4: (a) $^{13}$C ssNMR spectra of the indicated samples at 25 K, recorded with and without microwave (MW) irradiation to illustrate sensitivity enhancements from DNP. Spectra were acquired with the pulse sequence in Fig. S3b. Without MW irradiation, the strongest signals arise from natural-abundance $^{13}$C spins of frozen isopentane, which could not be separated completely from frozen protein solutions after rapid freezing. Signals from $^{13}$C-labeled protein molecules, but not from isopentane, are strongly enhanced by DNP. (b) Build-up of cross-polarized $^{13}$C ssNMR signals from U-$^{15}$N,$^{13}$C-FUS-LC (blue) and from residual GdmCl (orange) with increasing $\tau_{DNP}$ in measurements on the solution that was rapidly frozen from 4° C. Signal areas were measured from the regions indicated in the spectrum on the right and are normalized to their maximum values. Lines are fits with a single-exponential function, resulting in values of the build-up time $T_{DNP}$ in Table S1. The observation of significantly different $T_{DNP}$ values supports the existence of a phase-separated state in this frozen solution. (c) Comparison of DNP build-up curves for $^{1}$H signals and for cross-polarized $^{13}$C signals for frozen TYG-$^{15}$N,$^{13}$C-FUS-LC solutions. Signals were recorded with pulse sequences in Figs. S3a and S3b.

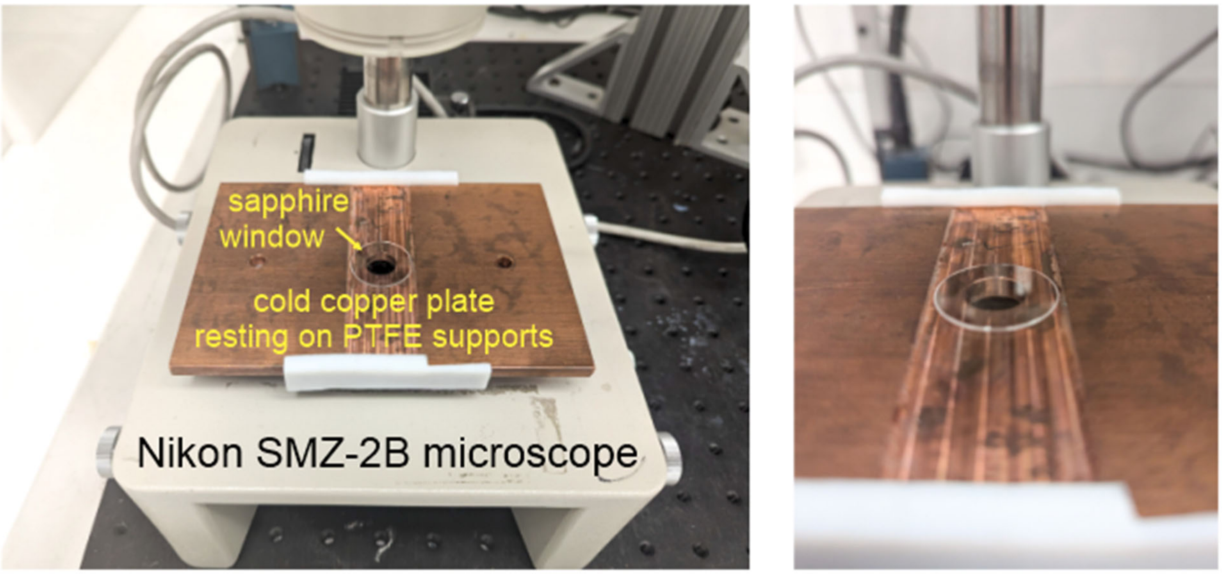


Figure S5: Apparatus for measuring optical microscope images of phase-separated FUS-LC droplets in the frozen state, as in Fig. 1g.

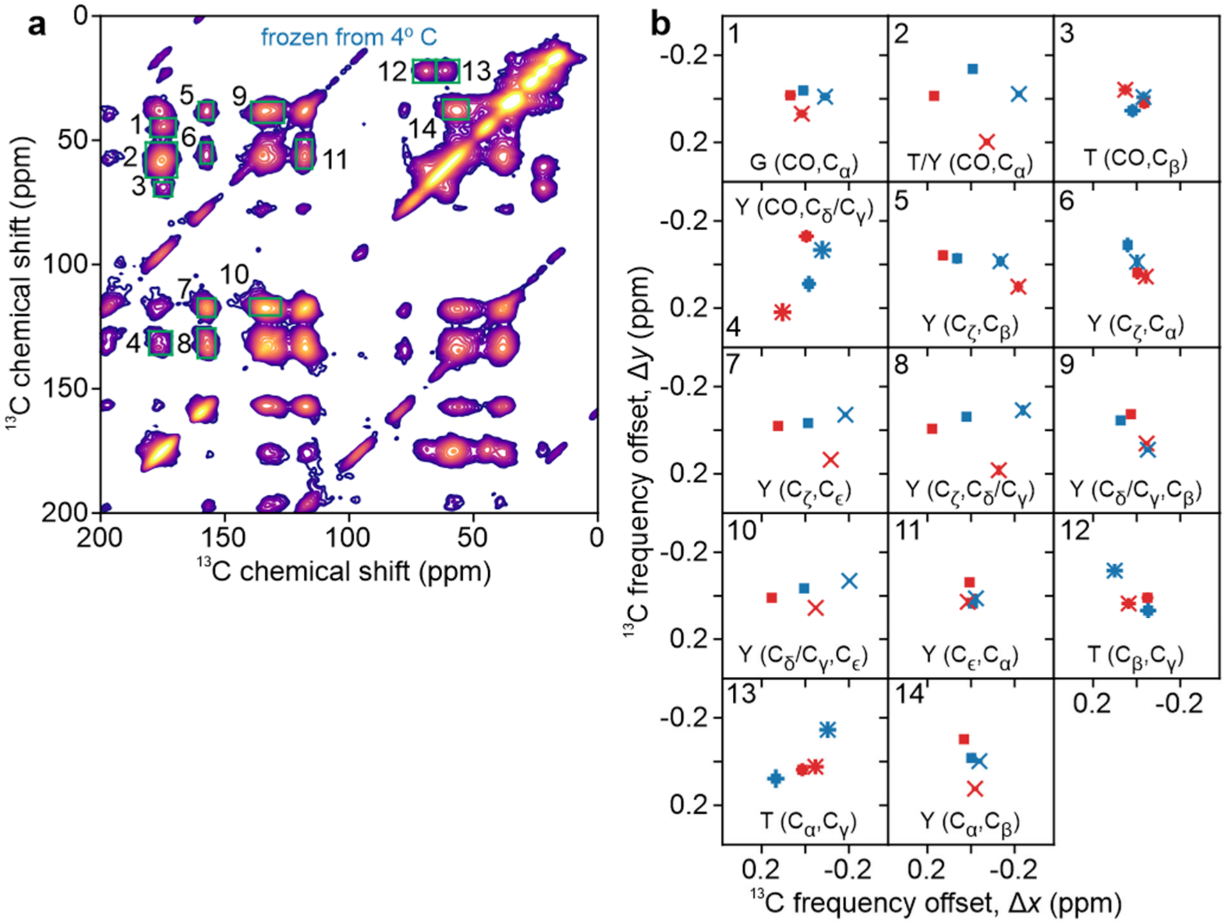


Figure S6: (a) DNP-enhanced 2D $^{13}$C-$^{13}$C ssNMR spectrum of a TYG-$^{15}$N,$^{13}$C-FUS-LC solution that was rapidly frozen 4° C (same as Fig. 3a). Center-of-mass values were calculated for crosspeak signals within green rectangles, as well as for crosspeaks related by symmetry across the diagonal of the 2D spectrum. (b) Center-of-mass values of the indicated crosspeaks, calculated for 2D spectra in Fig. 3a (blue) and Fig. 3b (red) and plotted relative to the average values. Values calculated for rectangular regions above and below the diagonal are shown with ■ and × symbols, respectively. Bars indicate uncertainties, calculated as explained in the main text.

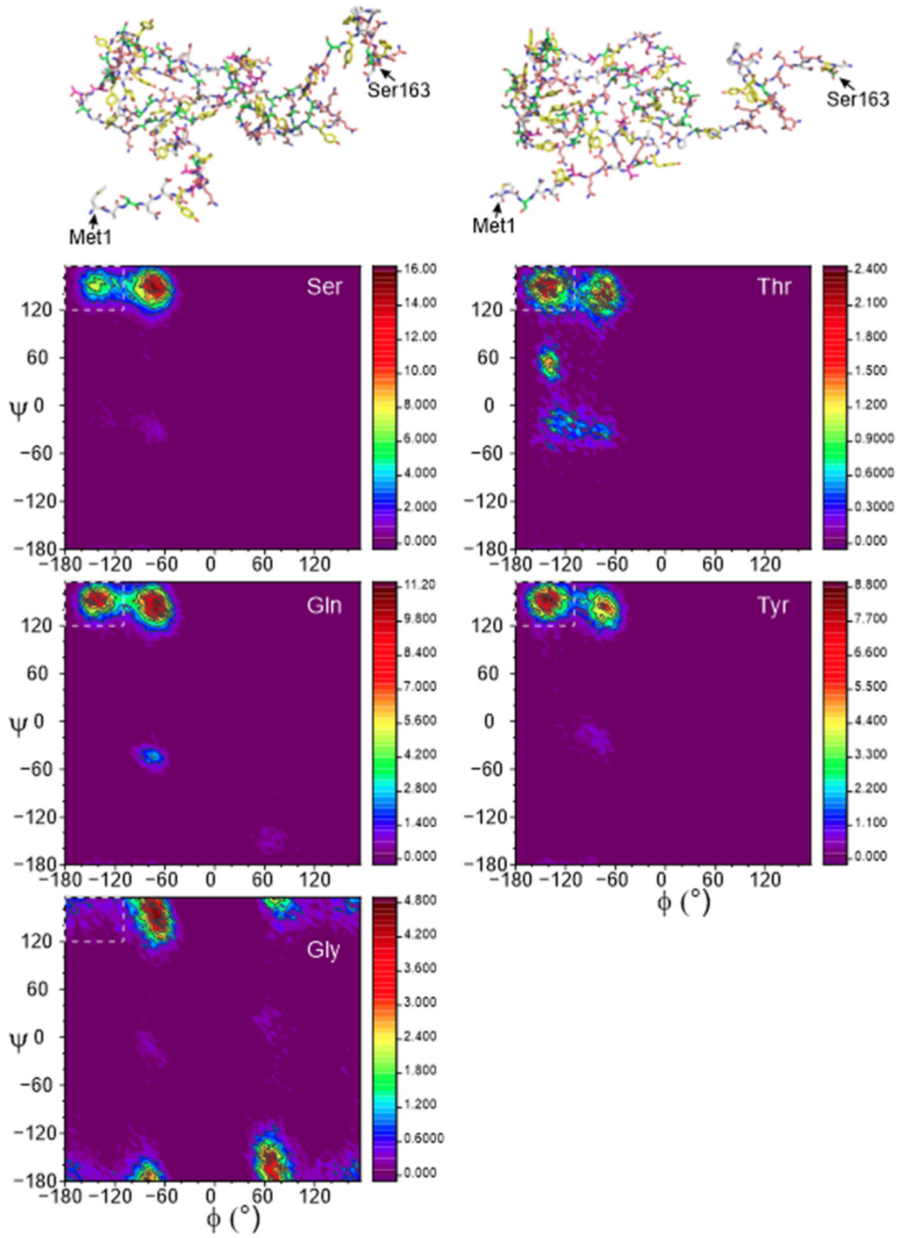


Figure S7: Plots of backbone ϕ,ψ torsion angle distributions (*i.e.*, Ramachandran plots) extracted from MD simulations on $FUS_{1\text{-}163}$. Configurations at the beginning and end of a 100 ns trajectory are shown at the top. These plots represent the combined distributions for all Ser, all Thr, all Gln, all Tyr, and all Gly residues in $FUS_{1\text{-}163}$. Torsion angles are binned in 5° increments. Regions enclosed by dashed white lines represent β-strand-like conformations.

Table S1: DNP enhancements of $^{13}C$ ssNMR signals ($\varepsilon_{DNP}$), and DNP build-up times ($T_{DNP}$) for FUS-LC and $FUS_{61-214}$ solutions that were rapidly frozen from temperatures above or below their phase-separation temperatures. Enhancement factors were measured for protein signals with $^{1}H$-$^{13}C$ cross-polarization (CP). Build-up times were measured for protein signals with CP, for $^{1}H$ signals, and for signals from natural-abundance $^{13}C$ of residual GdmCl detected with CP. Measurements on sel-$^{15}N$,$^{13}C$-$FUS_{61-214}$ used a lower-power microwave source for DNP than measurements on the FUS-LC samples, accounting for the smaller values of $\varepsilon_{DNP}$.

| Sample | Incubation temperature | $\varepsilon_{DNP}$ | $T_{DNP}$ (CP to protein) | $T_{DNP}$ ($^{1}H$ direct) | $T_{DNP}$ (CP to GdmCl) |
|---|---|---|---|---|---|
| U-$^{15}N$, $^{13}C$-FUS-LC | 50° C | 70 ± 6 | 2.6 s | - | 2.8 s |
| | 4° C | 54 ± 3 | 2.1 s | - | 3.3 s |
| TYG-$^{15}N$, $^{13}C$- FUS-LC | 50° C | 76 ± 6 | 5.2 s | 7.2 s | 10.5 s |
| | 4° C | 55 ± 4 | 2.6 s | 7.0 s | 8.6 s |
| sel-$^{15}N$,$^{13}C$-$FUS_{61-214}$ | 50° C | 36 ± 3 | 6.7 s | 6.7 s | - |
| | 4° C | 29 ± 2 | 8.3 s | 7.1 s | - |

Table S2: $^{13}C$ chemical shifts relative to sodium trimethylsilylpropanesulfonate (DSS) in ssNMR spectra of frozen U-$^{15}N$,$^{13}C$-FUS-LC solutions. Random coil values from Wishart *et al*. (ref. 83 of the main text) are given in parentheses. Error limits represent uncertainties in identifying positions of maximum intensity in 2D crosspeak signals. Values with large error limits correspond to sites with poorly resolved crosspeaks.

| Residue type | $^{13}C\alpha$ (ppm) | $^{13}CO$ (ppm) | $^{13}C\beta$ (ppm) | $^{13}C\gamma$ (ppm) | $^{13}C\delta$ (ppm) | $^{13}C\epsilon$ (ppm) | $^{13}C\zeta$ (ppm) |
|---|---|---|---|---|---|---|---|
| Gly | 45.0 ± 0.2 (45.1) | 174.8 ± 0.2 (174.9) | - | - | - | - | - |
| Gln | 55.5 ± 2.0 (55.7) | 176 ± 2 (176.0) | 31 ± 3 * (29.4) | 31 ± 3 * (33.7) | 180.4 ± 0.4 (180.5) | - | - |
| Thr | 61.0 ± 0.3 (61.8) | 175.1 ± 0.3 (174.7) | 69.2 ± 0.3 (61.8) | 22.3 ± 0.3 (21.5) | - | - | - |
| Tyr | 56.6 ± 0.4 (57.9) | 176.4 ± 0.4 (175.9) | 38.5 ± 0.4 (38.8) | 130.8 ± 0.4 (130.6) | 134.0 ± 0.4 (133.3) | 118.0 ± 0.4 (118.2) | 157.5 ± 0.4 (157.3) |
| Ser | 58 ± 2 (58.3) | 175 ± 2 (174.6) | 64 ± 2 (63.8) | - | - | - | - |

Table S3: $^{13}C$ chemical shifts relative to DSS in ssNMR spectra of frozen solutions of sel-$^{15}N$,$^{13}C$-FUS$_{61\text{-}214}$. Error limits represent uncertainties in identifying positions of maximum intensity in 2D crosspeak signals. Values with large error limits correspond to sites with poorly resolved crosspeaks.

| Residue | $^{13}C\alpha$ (ppm) | $^{13}CO$ (ppm) | $^{13}C\beta$ (ppm) | $^{13}C\gamma$ (ppm) | $^{13}C\delta$ (ppm) | $^{13}C\epsilon$ (ppm) | $^{13}C\zeta$ (ppm) |
|---|---|---|---|---|---|---|---|
| G65 | 45.0 ± 0.3 | 174.4 ± 0.3 | - | - | - | - | - |
| Q69 | 55 ± 2 | 177 ± 2 | 30 ± 2 | 33.8 ± 1.0 | 180.4 ± 0.4 | - | - |
| T78 | 61.1 ± 0.4 | 175.0 ± 0.4 | 69.5 ± 0.4 | 22.1 ± 0.4 | - | - | - |
| Y81 | 56.8 ± 0.3 | 176 ± 2 | 38.5 ± 0.3 | 131 ± 2 | 134.1 ± 0.3 | 118.0 ± 0.3 | 157.5 ± 0.3 |
| S86 | 58 ± 2 | 175 ± 2 | 64 ± 2 | - | - | - | - |
